# Conditioned versus innate effort-based tasks reveal divergence in antidepressant effect on motivational state in male mice

**DOI:** 10.1101/2025.02.04.636449

**Authors:** Foteini Xeni, Caterina Marangoni, Lynn Lin, Emma S J Robinson, Megan G Jackson

## Abstract

Antidepressant-induced apathy syndrome is reported in a high number of patients. It is characterised by loss of motivation for daily activities and emotional blunting. It has a negative impact on quality of life and treatment outcome, yet the changes in underlying neurobiology driving this syndrome remain unclear. To begin to address this, a comprehensive understanding of how different classes of antidepressant treatment impact on behaviours relevant to apathy is critical. Rodent motivation for reward is commonly assessed using effort-based operant conditioning paradigms such as the Effort for Reward task. However, motivation to perform spontaneous/innate behaviours may provide additional insight into changes in behaviour reflective of daily activities. We tested the acute and chronic effects of antidepressants on the Effort for Reward task, and the spontaneous/innate Effort Based Forage task. Acute treatment revealed important divergence in drug effect between tasks, where selective serotonin reuptake inhibitor (SSRI)/serotonin and noradrenaline reuptake inhibitor (SNRI) treatment impaired foraging behaviour in the Effort Based Forage task, but enhanced high effort, high value reward responding in the Effort for Reward task. Treatment with a noradrenaline reuptake inhibitor (NRI) or multimodal agent impaired foraging behaviour but did not affect high reward responding in the Effort for Reward task. Conversely, chronic treatment with an SSRI but not SNRI enhanced motivated foraging behaviour but led to a general impairment in Effort for Reward task output. Together, these data demonstrate that SSRI treatment induces opposing effects on conditioned versus innate motivation which may have significant translational relevance when interpreting drug effect.

## Introduction

In the UK alone, 1 in 6 people were reportedly taking antidepressants in 2022-23, and the number is projected to rise further still (1). While patients do report relief from negative emotional symptoms, a large proportion of patients report the inability to experience a full range of emotions, alongside a profound deficit in their motivational state following treatment. This clinically significant state termed selective serotonin reuptake inhibitor (SSRI) or antidepressant-induced apathy syndrome results in a negative impact on both quality of life and treatment outcome. Prevalence of apathy syndrome is dependent on antidepressant class, occurring in 20-90% patients taking SSRIs and 5.8-50% of patients taking antidepressants including selective serotonin and noradrenaline reuptake inhibitors (SNRIs)(2). A task probing the effects of SSRI treatment found a blunted responsiveness during reinforcement learning and decision-making in human participants demonstrating the impact of these residual symptoms on reward-related cognition (3). Treatment with SSRIs reduces activation in brain reward networks in human volunteers early in the treatment regime (4). However, despite its scientific and clinical importance, the underlying neurobiology of antidepressant-induced apathy remains unknown. To address this, the use of a translational animal model to probe changes relevant to motivational deficit and emotional blunting induced by antidepressant administration at the brain and behavioral level is important.

The Effort for Reward task (EfR) is an operant-based method for studying motivated behaviour in rodents (5) and has been reverse translated in human participants (6). In this task, the rodent is food restricted, and trained to make an effortful number of operant responses to receive a palatable food reward, or consume a lower value but freely available standard lab chow. The rodent’s motivational state is captured by its willingness to exert effort to receive high value reward. The Effort-Based Forage task (EBF) is an ethological, spontaneous measure of motivational state based on the intrinsic drive of the mouse to forage nesting material under various effortful and environmental contingencies (7). By bypassing the requirement for food restriction or conditioned learning, this task captures self-initiated or intrinsically rewarding behaviours, where the effort applied to foraging is innately rewarding.

While both tasks show sensitivity to dopaminergic modulation (5, 7–9) and are unified under the umbrella of ‘motivation tasks’, they may capture different aspects of motivational processing and associated neurobiological mechanisms. These differences may have important translational implications. A recent study found distinct neural alterations in learned reward processing (monetary) versus natural (intrinsically rewarding) rewards (e.g., a favourite song) in the context of MDD (10). Given that natural rewards pervade our daily lives, it is important to ask the question: how do antidepressants impact on these dimensions of motivated behaviour relevant to apathy behaviour?

Here, we tested a range of commonly prescribed SSRI and non-SSRI antidepressants to capture the effects of a diverse range of binding profiles and mechanisms of action within both the EfR and EBF tasks. We utilised animal-equivalent, clinically-relevant doses and voluntary oral administration to translate to clinical conditions. Acute treatment was tested first followed by chronic treatment of two antidepressants representative of the SSRI and SNRI classes to capture short and longer term effects of the drug on behaviour. Based on delayed onset of therapeutic efficacy we predicted that chronic but not acute treatment would impair motivated behaviour. Based on differences in prevalence of apathy between antidepressant classes we additionally predicted non-SSRIs may affect motivation less than SSRIs.

## Methods

### Subjects

4 cohorts of male C57bl/6J mice (Envigo UK) were used in these experiments (see **S1** for summary). Mice were singly housed in open-top Techniplast 1284 conventional caging to minimise psychosocial stress associated with aggression (11). Mice underwent twice daily health checks to ensure good welfare throughout testing. Mice were kept in temperature-controlled conditions (∼21 °C) and a 12:12h reversed light-dark cycle. Animals were habituated to handling and managed following the principles of the 3Hs framework (12, 13). Further husbandry details are provided in (**S2**). Mice underwent testing during their active phase. Standard laboratory chow (Purina, UK) and water were provided *ad libitum*, apart from during operant training and testing, when mice were mildly food restricted. Food was removed ∼4h before testing and replaced 1h after testing. During food restriction, weights were monitored at least once a week and maintained to at least 85% of their free feeding weight relative to their normal growth curve. Sample sizes were based on detecting a large effect size (∼1.0) with mean difference and variance based on previous behavioural studies using similar tasks and acute/chronic pharmacological manipulations (7, 9). In all experiments the experimenter was blind to treatment. In the acute studies, each mouse received each dose and vehicle in a fully randomised, within-subject counterbalanced Latin square-design, over four test sessions. While for the chronic studies a between-subject design was utilised. All experiments were conducted in accordance with the Animals Scientific Procedures Act (United Kingdom, 1986) and were approved by the University of Bristol Animal Welfare and Ethical Review Body (AWERB). This study was pre-registered in the Open Science Framework at https://osf.io/ny37m/.

### Operant training

Operant training was consistent with the protocol reported in (9). Mice were trained in sound-proofed operant boxes (MedAssociates) run on Klimbic software. Each operant box consisted of 2 nose poke apertures and a centrally located food magazine connected to a reward pellet dispenser (20mg rodent tablet, TestDiet, Sandown). A Lifecam HD 3000 webcam was positioned in the ceiling of each box, with a view of the interior of the operant box (**fig.1A**).

**Figure 1.**
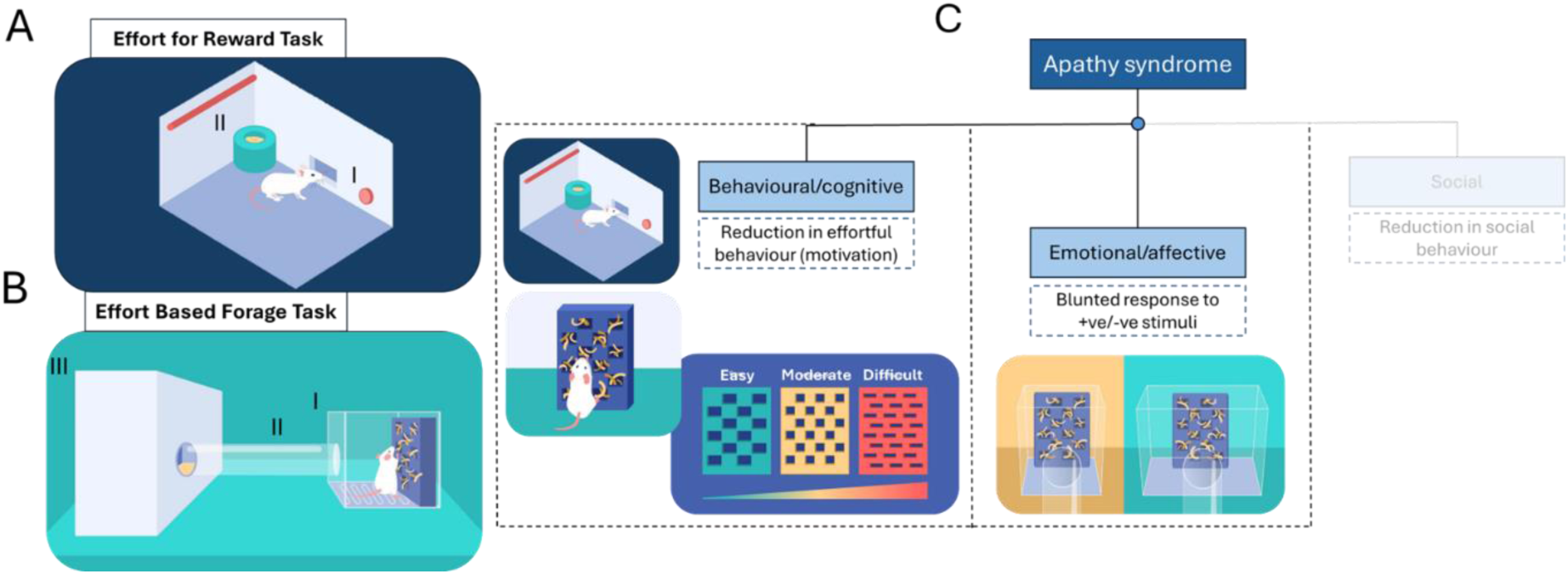
Overview of the Effort for Reward Task and Effort-Based Forage Task and their mapping to apathy behavioural dimensions. **A** In the EfR task, the mouse is trained in an operant box and is given the option between responding at the active nose poke four times (I) to receive a high value reward, or can consume freely available standard powdered chow placed on the other side of the magazine (II). The number of high effort trials the mouse completes gives a readout of motivational state. **B** In the EBF task, the mouse is placed in a safe and enclosed home area (III) with free access to food and water. Throughout the session, the mouse can choose to traverse the tube (II) connecting to a more open forage area (barred lid to allow air flow and use of clear Perspex) (I). A nesting box is placed in the forage area, and the mouse can choose to effortfully forage nesting material via apertures in the face plate of the nesting box. The mouse can then choose to pick up the nesting material and shuttle it back to the home area (III). **C** Apathy has three core dimensions including a behavioural/cognitive domain and an emotional/affective domain (19). The dashed boxes below denote how behaviours change clinically within these domains. Reduction in effortful behaviour can be measured by the Effort for Reward Task and standard Effort Based Foraging Task. A variant of the Effort-Based Foraging Task called the effort curve paradigm can test effort-based modulation of behaviour by changing effort contingency by altering aperture size in the nesting box faceplate. The emotional/affective domain of apathy can be tested by assessing foraging behaviour in a more open (aversive environment). A blunting in affective response to a more aversive environment may be indicative of emotional blunting (7, 20). Social motivation is not yet tested in these paradigms.

The operant training was completed in 3 stages outlined in **S3**. In the final stage of training, the mice progressed through FR1, 2 and 4, where the number corresponds to the number of nose pokes at the active aperture necessary for the delivery of a reward pellet. Mice completed each stage on an individual basis following 2 consecutive sessions of 30+ trials. A trial refers to the delivery of a reward following the required number of nose pokes. Testing began once training was complete.

### Effort for Reward task

A test session consisted of a 30 minute FR4 session with the addition of a bowl of powdered standard chow positioned in front of the inactive nose poke aperture (**fig.1A**). The chow is accessed via a 1.27cm hole in the bowl lid to facilitate chow access but prevent digging. The FR4 requirement to obtain a single reward pellet represents the high effort-high value reward option, while the freely available chow represents the low effort-low value reward option. Throughout the session, the mouse can freely choose between these options. The mice completed five test sessions prior to pharmacological testing to habituate to the presence of the chow bowl. Videoed behaviour at the chow bowl was additionally scored using an in-house developed application (9). The primary outputs of the task were number of high effort, high value trials completed and amount of chow consumed (g). Secondary bowl outputs are summarised in **S4**.

#### Acute effects of SSRI and non-SSRI antidepressants

N = 16 mice were used for these experiments. Baseline FR4 sessions were conducted on Monday and Thursday, while test sessions were conducted on Tuesday and Friday (**S5**) to ensure drug washout between test sessions. Mice were administered the drugs orally in a palatable solution with a dose volume 10ml/kg and were trained to drink voluntarily from a syringe as previously described (7, 12). The following drugs were tested: escitalopram (1, 3, 10mg/kg, t-60mins, Lundbeck ltd), citalopram (1, 3, 10mg/kg, t-60mins, HelloBio), fluoxetine (1, 3, 10mg/kg, t-4hours, HelloBio), sertraline (1, 3, 10mg/kg, t-4hours, HelloBio), venlafaxine (1, 3, 10mg/kg, t-60mins, Rosemont), reboxetine (0,3, 1, 3mg/kg, t-30mins, Sigma), and vortioxetine (0.3, 1, 3mg/kg, t-2hours, MedChem Express). Drug vehicles are reported in **S6**. Dose ranges were based on human dose ranges using the animal equivalent dose equation (14) and were aligned with clinically relevant doses where possible (**S6**). Pretreatment times (t-) were based on T_max_.

#### Effects of chronic treatment with escitalopram (SSRI) and venlafaxine (non-SSRI)

Vehicle was administered to half the mice (n=12), while the other half (n=12) received escitalopram 3mg/kg (1st cohort) or venlafaxine 10mg/kg (2nd cohort). Doses were based on findings from acute treatment. Dosing occurred 14 days prior to the experiment (allowing time to steady state concentration and relevant receptor adaptation including 5HT-1A desensitisation (15–18)) and continued during testing. The drug was administered once per day at ∼2pm. Mice underwent four baseline FR4 sessions preceding a test session (**S5**). Sessions preceded dose time so that acute effects of the drug did not impact performance.

### Effort-Based Foraging task

#### Habituation and output

N=16 mice were first habituated to the EBF arena across three sessions as previously described (7) and outlined in **S7**. The final stage of habituation consisted of an initial 4hr foraging session where a custom-designed nesting box ((7), https://github.com/meganjackson13/Bedding-box-3D-files) was placed in the foraging area filled with 18g of Sizzlenest (Datesand, UK), which could be accessed via 1.5 cm^2^ apertures (lowest level effort). Arena details are provided in **S8A-C** and **fig.1B**.

The nesting box is weighed pre- and post-session providing a measure of total nesting material foraged (pulled from nesting box). Amount of nesting material pulled from the nesting box but left on top of the barred floor of the forage arena was also weighed. The two main output measures were 1) *total nesting material shuttled to the home area (g)* which reflects the amount of nesting material pulled from the nesting box (foraged) and carried through the tube to the home area and 2) *% nesting material foraged taken to the home area* which reflects of the total bedding foraged, how much is shuttled to the home area instead of left in the open, expressed as a percentage. A table of output measures are provided in **S9**.

#### Acute effects of SSRI and non-SSRI antidepressants

Post-habituation mice underwent three 2-hr sessions using the 1cm^2^ apertures (moderate effort), to ensure stability in performance before starting drug testing. Each test session was separated by two days to ensure drug washout between sessions (**S10**). Mice were administered the drugs consistent with the EfR studies above. General activity in the forage area was recorded using an in-house passive infrared system (**S11**)(21). Where task activity changes were detected, mice underwent home-cage activity monitoring to test for general impairments in locomotion not specific to the task environment (**S12**).

#### Effects of chronic treatment with escitalopram and venlafaxine

Following testing in the EfR task, mice were habituated to the EBF task as described. To test effort-based modulation of behaviour, mice underwent effort curve testing. In this paradigm the effort required for foraging is modulating by changing the aperture sizes within the faceplate. Mice were presented with each of these faceplates in a within-subject counterbalanced design, across three test sessions (**S8**).

Mice then underwent two 2-h foraging sessions using the 1.5 cm^2^ aperture where the nesting box was placed in either an enlarged or standard forage area, counterbalanced. This was to test affective reactivity driven by mice finding more open, novel spaces aversive (22)(**fig.1C**).

### Statistics

Data were tested for normality using a Shapiro-Wilk test. Where data were normally distributed a repeated measure one-way ANOVA (dose response) or independent t-test (between-subject vehicle versus treatment) were performed where appropriate. Where a main effect was observed (p<0.05), Dunnett’s post-hoc comparisons were carried out. Where data violated normality, the Friedman test with Dunn’s post-hoc analysis or the Mann-Whitney U test was conducted. Where two-factor analysis was required (treatment*effort, treatment*arena size), a RM two-way ANOVA with Sidak’s multiple comparison was carried out. Trend-level main effects (p<0.1) were reported but not further analysed. Outliers were defined as values 2 standard deviations from the mean. For each measure, outliers were replaced with the group mean to permit repeated measure analysis. A table summarising exclusions is provided in **S13**. Data analysis and graphing were performed using GraphPad Prism v10.0.3.

## Results

### Acute SSRI treatment exerts opposing effects in the Effort for Reward and Effort-Based foraging task

Escitalopram reduced nesting material shuttled (Q=29.00, p<0.0001, (3mg/kg, p=0.0024), (10mg/kg, p<0.0001)) in the EBF task (**Fig.2A**). Escitalopram also reduced % shuttled (F_(2.009,30.14)_=6.045, p=0.0061, (3mg/kg, p=0.0186), (10mg/kg, p=0.0238) (**Fig.2B**). Notably 3 and 10mg/kg reduced general activity in the forage area (p=0.0057, p=0.0064, respectively) (**S14**). However, there was no effect on home cage activity (**S15**). Escitalopram increased trials completed in the EfR task (F_(1.997,29.96)_=11.68, p=0.0002, (1mg/kg, p=0.0646), (3mg/kg, p=0.0003), (10mg/kg, p=0.0044)) (**Fig.2C**). There was no effect of escitalopram on chow consumed (**Fig.2D**). 10mg/kg reduced chow bowl bouts (p=0.0032), but no other secondary bowl output measure was affected (**S16**). Citalopram showed a similar pattern of results in both tasks (**S17**).

**Fig. 2.**
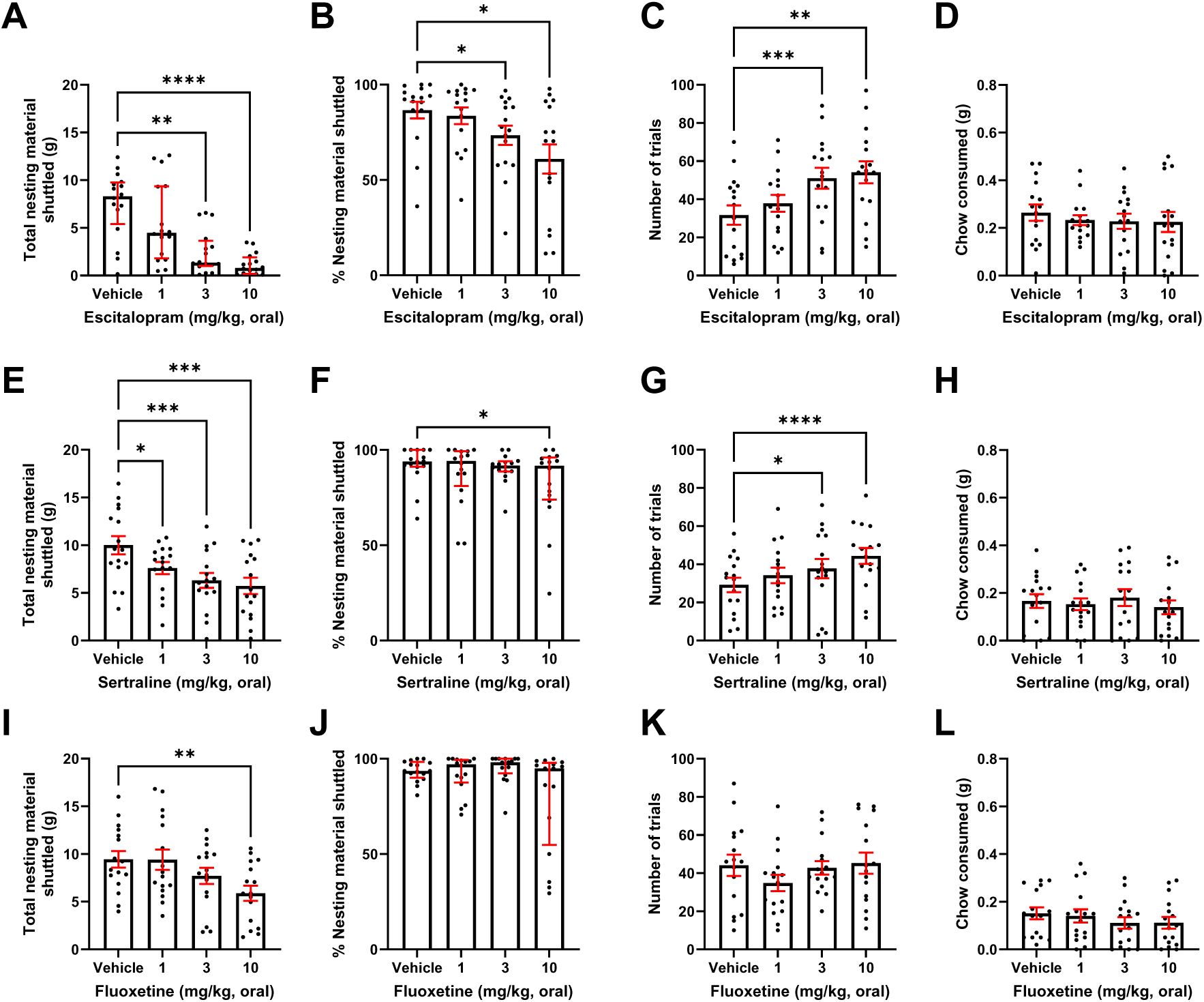
Acute SSRI treatment exerts opposing effects in the EfR and EBF task. Mice were dosed with a range of SSRIs and underwent the EBF or the EfR task. **A** 3 and 10mg/kg escitalopram caused a reduction both in the total nesting material shuttled (p< 0.001), and **B** on % of nesting material shuttled (p<0.05). **C** 3 and 10mg/kg escitalopram increased number of high-effort, high-value trials performed in the EfR task (p<0.001, p<0.01) **D** with no effect on chow consumed. **E** Sertraline reduced total nesting material shuttled in at all doses (p<0.05, p<0.001), and **F** 10mg/kg sertraline reduced the % of nesting material shuttled (p<0.05). **G** 3 and 10mg/kg sertraline increased the number of high-effort, high-value trials performed in the EfR task (p<0.05, p<0.0001) with **H** no effect on chow consumed. **I** 10mg/kg fluoxetine reduced the total amount of nesting shuttled (p<0.01), with **J** no effect on % nesting material shuttled **K**. Fluoxetine had no effect on high-effort trials and **L** on chow consumed in the EfR task. Error bars are mean ±SEM, one-way ANOVA was performed to detect drugs effect. For non-parametric distributions, the median and interquartile range were used, *p<0.05, **p<0.01, ***p<0.001, ****p<0.0001.

Sertraline decreased nesting material shuttled (F_(2.990,44.86)_=12.06, p<0.0001, (1mg/kg, p=0.0197), (3mg/kg, p=0.0008), (10mg/kg, p=0.0002)) (**Fig.2E**) and decreased % shuttled (Q=8.059, p = 0.0448, (10mg/kg, p=0.0185)) (**Fig.2F**). Forage area activity was not affected (**S14**). Sertraline increased number of trials (F_(2.394,35.91)_=11.82, p<0.0001, (1mg/kg, p=0.0844), (3mg/kg, p=0.0400), (10mg/kg, p<0.0001)) (**Fig.2G**). Chow consumed (**Fig.2H**) and secondary parameters were not affected (**S16**).

Fluoxetine decreased nesting material shuttled (F_(2.475,37,13)_=5.797, p=0.0039, (10mg/kg, p=0.0060)) (**Fig.2I**), and % shuttled (Q=9.627, p=0.0220) however post-hoc analysis did not reach significance (**Fig.2J**). Forage area activity was not affected (p>0.05) (**S14**). Fluoxetine trended towards a decrease in number trials (F_(2.763,41.44)_=2.990, p=0.0456, (1mg/kg, p=0.0590)) (**Fig.2K**), but did not affect chow consumed (**Fig.2L**). There was no change in all other EfR task parameters (**S16**).

### EBF task shows greater sensitivity to acute treatment with non-SSRI antidepressants

Venlafaxine reduced total nesting material shuttled (F_(2.618,39.27)_=7.114, p=0.0010, (3mg/kg, p=0.0346), (10mg/kg, p=0.0031)) (**Fig.3A**) and % shuttled (Q=12.73, p=0.0053) however post-hoc analysis did not reach significance (**Fig.3B**). Forage area activity was not affected (**S14**). Venlafaxine increased trials (F_(2.010, 29.47)_=4.897, p=0.0145, (10mg/kg, p=0.0041)) (**Fig.3C**). There was no effect on chow consumed or secondary output measures (**Fig.3D**, **S18**).

**Fig.3.**
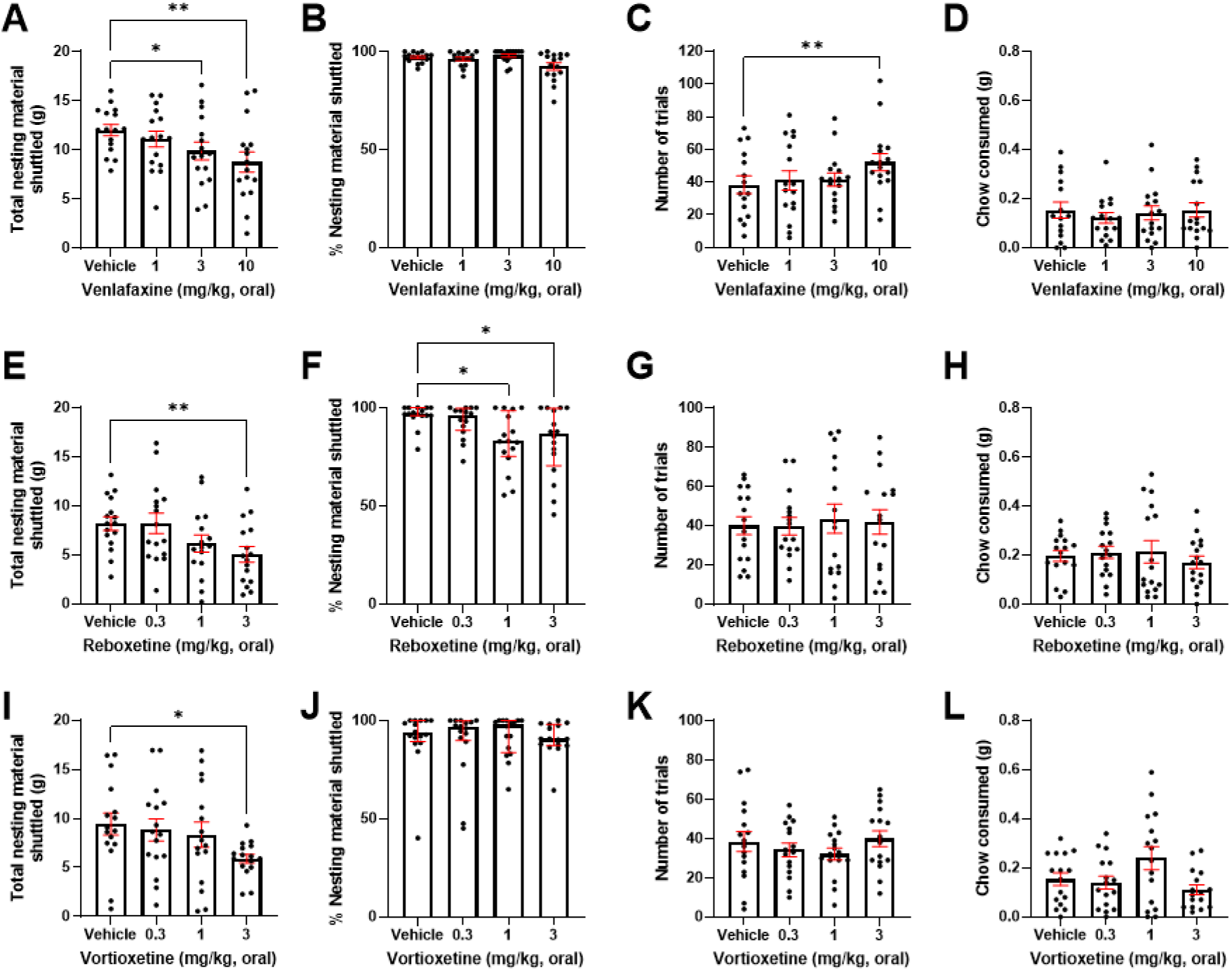
EBF task shows greater sensitivity to acute treatment with non-SSRI antidepressants compared to the EfR task. Mice were dosed with a range of non-SSRI antidepressants and underwent the EBF or the EfR task **A**. 3 and 10mg/kg venlafaxine caused a reduction in the total nesting material shuttled (p<0.05, p<0.01) with no effect on **B** the % of nesting material shuttled **C.** 10mg/kg venlafaxine increased the number of completed high-effort trials (p<0.01), with no effect on **D** chow consumed in the EfR task. **E.** 3mg/kg reboxetine reduced total nesting material shuttled (p<0.01), and **F**. 1 and 3mg/kg reboxetine caused a reduction on the % of nesting material shuttled (p<0.05) **G.** Reboxetine had no effect on number of trials and **H** chow consumed **I.** 3mg/kg vortioxetine reduced total nesting material shuttled (p<0.05). Vortioxetine had no effect on **J** % of nesting material shuttled or **K** number of trials and **L** chow consumed in the EfR task. Error bars are mean ±SEM, one-way ANOVA was performed to detect drugs effect, for non-parametric distributions, the median and interquartile range were used *p<0.05, **p<0.01.

Reboxetine reduced nesting material shuttled (F_(2.187,32.81)_=4.742, p=0.0134, (3mg/kg, p=0.0019)) (**Fig.3E**) and % shuttled (Q=12.11, p=0.0070, (1mg/kg, p=0.0278), (3mg/kg, p=0.0339) (**Fig.3F**). Reboxetine had no effect on forage area activity (**S14**). It had no effect on primary EfR task parameters (**Fig.3G&H**) but increased average duration at the chow bowl (3mg/kg, p=0.0219) and trended towards increased number of bouts at the chow bowl (p=0.0851) (**S18**).

Vortioxetine reduced nesting material shuttled (F_(2.759,41.39)_=3.235, p=0.0352, (3mg/kg, p=0.0478)) (**Fig.3I**) but did not affect % shuttled (**Fig.3J**), or forage area activity (**S14**). It had no effect on number of trials completed in the EfR task (p>0.05) (**Fig.3K**) but trended toward a decrease in chow consumed (F_(1.627,24.40)_=6.160, p=0.0100, (3mg/kg, p=0.0587)) (**Fig.3L**). There was no change in secondary task parameters (**S18**).

### Chronic escitalopram treatment increased foraging behaviour but induced general impairment in the EfR task

There was no effect of treatment on initial foraging during habituation (**S19**). When assessed using the effort curve paradigm there was a main effect of treatment (F(_1,20_)=6.484, p=0.0192) where treatment increased nesting material shuttled, but no aperture size*drug interaction (**Fig.4A**). Post-hoc comparisons didn’t reach significance. There was no effect of aperture on % shuttled, although there was a trend towards aperture size*drug interaction (F(_2,40_)=2.499, p=0.0949) (**Fig.4B**).

**Fig.4.**
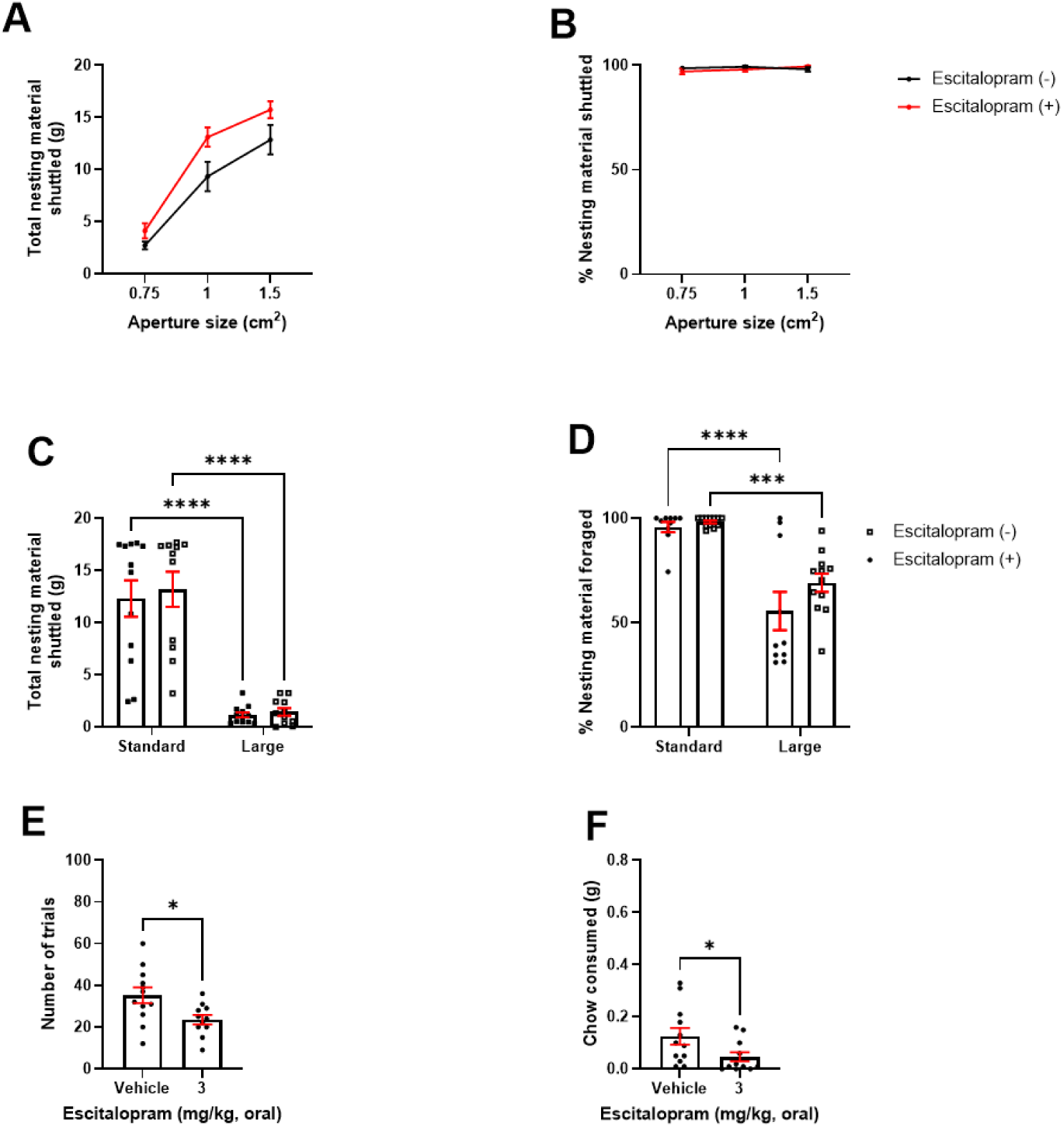
Chronic escitalopram treatment improved foraging behaviour under varying effort contingencies but induced general impairment in the EfR task. Mice were dosed with 3mg/kg escitalopram for 2 weeks and underwent behavioural testing. **A** Increasing aperture sizes increased total nesting material shuttled in both groups. Escitalopram induced a general increase in nesting material shuttled. **B** There was no difference in % nesting material shuttled at different apertures**. C** Both groups showed reduced total nesting material shuttled in a larger forage area (p<0.0001)**. D** % nesting material shuttled was reduced in the larger forage area in both groups (p<0.0001 (vehicle), p<0.001 (treated)). **E.** Treated mice showed a reduction in high-effort, high reward selection option (p<0.05) and a reduction in **F** chow consumed (p<0.05). Error bars are mean ±SEM, unpaired t-test or RM two-way ANOVA was performed to detect the drug effect. For non-parametric distributions, the median and interquartile range were used *p<0.05, ***p<0.001, ****p<0.0001.

A larger forage area reduced total nesting material shuttled in both groups (F(_1, 21_)=107.5, p<0.0001, (vehicle and 3mg/kg, p<0.0001)) however there was no effect of treatment, or size*treatment interaction (**Fig.4C**). A larger forage area also reduced % shuttled (F(_1,20_)=55.98, p=0.0001, (vehicle, p<0.0001), (3mg/kg, p=0.0003)), but there was no effect of treatment or size*treatment interaction (**Fig.4D**). Treated mice completed less trials (t(_21_)=2.588, p=0.0172) and consumed less chow (t(_22_)=2.164, p=0.0416) (**Fig.4E&F**). Treated mice spent less time at the chow bowl (t(_22_)=2.259, p=0.0346) but no other secondary output measure was affected (**S20**).

### Chronic venlafaxine treatment showed limited effect in either the EBF or EfR tasks

There was no effect of chronic venlafaxine treatment on initial foraging during habituation (**S19**). When assessed using the effort curve paradigm there was no treatment or aperture size*treatment interaction on total or % nesting material shuttled (**Fig.5A&B**). A larger forage area reduced total nesting material shuttled in both groups (F(_1, 21_)=1478, p<0.0001, vehicle and 3mg/kg, p<0.0001). There was a size*treatment interaction (F(_1, 21_)=8.266, p=0.0091), but no effect of treatment. In the enlarged forage area only, venlafaxine-treated mice trended towards shuttling more nesting material than non-treated mice (p=0.0536) (**Fig.5C**). A larger forage area also reduced % shuttled in both groups (F(_1,21_)=69.93, p<0.0001, (vehicle, p<0.0001), (3mg/kg, p=0.0002). There was no size*treatment interaction but there was a trend level effect of treatment (F(_1,21_)=3.353, p=0.0813) (**Fig.5D**). There was no effect of chronic venlafaxine treatment in the EfR task (**Fig.5E&F,S19**).

**Fig.5.**
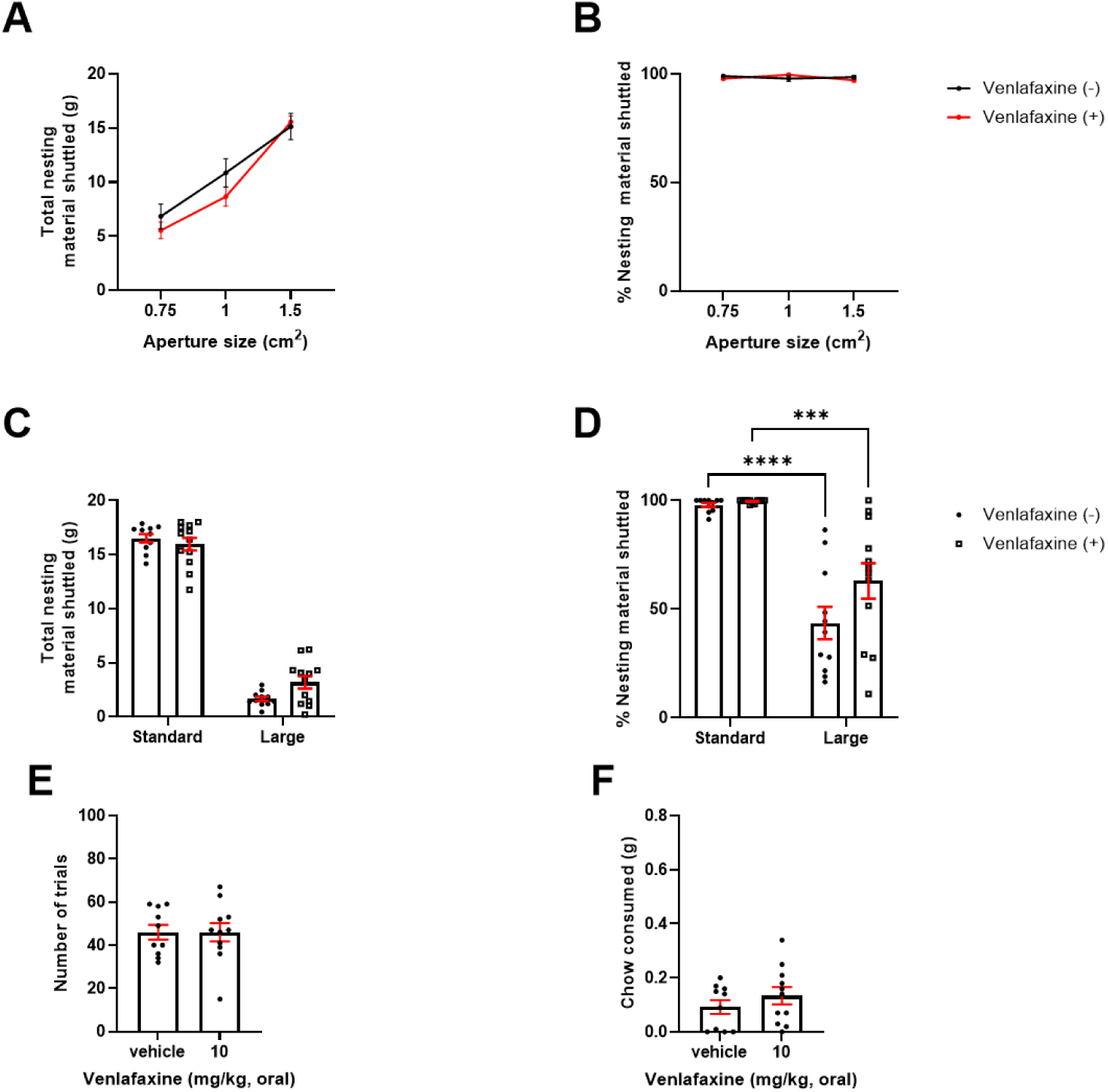
Chronic venlafaxine treatment showed limited effect in either the EBF or EfR tasks. Mice were dosed with 10mg/kg venlafaxine for 2 weeks and underwent behavioural testing. **A** Increasing aperture sizes increased total nesting material shuttled in both groups. There was no effect of treatment. **B** There was no difference on % nesting material shuttled at different apertures. **C** Both groups showed reduced total nesting material shuttled in a larger forage area (p<0.0001 (vehicle), p<0.001 (treated group)). **D** % nesting material shuttled was reduced in the larger forage area in both groups (p < 0.0001). **E** Treated mice showed no effect on number of trials completed and **F** chow consumed in the EfR task (p > 0.05). Error bars are mean ±SEM, unpaired t-test or RM two-way ANOVA was performed to detect drug effect, *p<0.05, ***p<0.001, ****p<0.0001. For non-parametric distributions, the median and interquartile range were used.

## Discussion

Comparison of the effect of acute antidepressant treatment in the Effort-Based Forage (EBF) task and Effort for Reward (EfR) task revealed an interesting divergence in behavioural effect. Acute treatment with escitalopram, citalopram, sertraline and venlafaxine dose-dependently reduced nesting material shuttled in the EBF task. However, these drugs increased the number of high effort, high value trials completed in the EfR task. The highest dose of fluoxetine, reboxetine and vortioxetine reduced nesting material shuttled in the EBF task but did not affect EfR task performance. Escitalopram, citalopram, sertraline and reboxetine also impaired % nesting material shuttled. In contrast to acute effects with escitalopram, chronic treatment induced a general increase in nesting material shuttled, and an overall reduction in EfR task performance. Chronic treatment with venlafaxine had no effect on EBF or EfR task performance. The following discussion will consider how these findings contribute to our understanding of the effects of antidepressant drugs on apathy-related behaviour.

### EBF performance is impaired by acute SSRI and non-SSRI treatment, while EfR performance is increased with SSRI treatment only

Acute treatment may provide insight into initial neurobiological and behavioural changes that have relevance to longer-term effects. Notably, all tested SSRIs acutely decreased nesting material shuttled in the EBF task, indicating that they induce an immediate reduction in intrinsically motivated behaviour. These findings were unlikely driven by a general impairment in locomotor behaviour or enhanced aversion to the forage area, as activity levels in the forage area/home cage remained unchanged. With the exception of fluoxetine, the SSRIs additionally reduced % nesting material shuttled. Previous work showed that in the standard arena format, aged but not corticosterone-treated mice showed a reduction in % nesting material shuttled, suggesting transportation of nesting material may reflect an additional effortful component of foraging behaviour. Previous work found this measure was unaffected by dopaminergic modulation (7), suggesting this measure is sensitive to differences between dopaminergic and serotonergic-mediated changes that require further investigation.

In contrast, and with the exception of fluoxetine, we observed increased effortful conditioned responding in the EfR task with SSRI treatment, indicating instead, an increase in motivated behaviour. Differences in drug effect between tasks suggests that while both capture aspects of motivated behaviour, there may be translationally-relevant differences in their underlying neurobiological mechanisms. EfR task performance is highly sensitive to dopaminergic modulation, where an increase in mesolimbic dopamine increases high effort, high value reward responding (5, 8, 9). The positive acute effects of SSRIs in the EfR task could be associated with the ability of 5-HT to modulate mesolimbic DA neurons. Previous work has shown that low-dose escitalopram increased the firing rate and burst firing of DA neurons in the ventral tegumental area (VTA) (23). Selective stimulation of VTA-projecting dorsal raphe nucleus 5-HT neurons strongly reinforced nose-poke self-stimulation behaviour in mice (24). Other work found acute fluoxetine or escitalopram treatment impaired high effort trial performance in conditioned effort-based decision-making tasks (25–27). However, use of intraperitoneal administration combined with choice of higher doses in these studies will engage higher levels of receptor occupancy, making comparison difficult.

Innate foraging behaviour measured by the EBF task may engage regions beyond simple bidirectional modulation of the dopaminergic mesolimbic circuitry. Innate behaviours including nest building are mediated by the prefrontal cortex (PFC)-mediated circuits (28). Elevated 5-HT may reduce glutamatergic activity in the PFC via direct or indirect inhibition of pyramidal neurons, which may suppress foraging behaviour (29). Gaining further insight into differences in mechanism between tasks could facilitate our understanding of their translation to motivational deficit induced by apathy syndrome.

Use of non-SSRI antidepressants may provide further insight into how distinct antidepressant mechanisms can differentially modulate behavioural output. The SNRI venlafaxine showed a similar profile of behavioural effects to the SSRIs, suggesting the serotonergic component of the drug may modulate the behavioural differences between tasks. Indeed, reboxetine, an RNI, did not affect performance in the EfR task. Vortioxetine also did not impact EfR performance. While vortioxetine does act as a SSRI, it has a unique binding profile with antagonistic/agonistic properties at a range of different 5-HT receptors (30) which may negate the acute behavioural effects observed with the other serotonergic drugs. Both reboxetine and vortioxetine reduced nesting material shuttled in the EBF at the highest dose. Impairment in the EBF task at a behaviourally inert dose in the EfR task suggests that the EBF task is particularly sensitive to neurochemical changes.

Together, these data suggest that intrinsically motivated behaviour is impaired by acute oral SSRI treatment. However, learned/conditioned motivated behaviours appear to be enhanced. Fluoxetine’s divergence from the rest of its class may relate to its reduced binding specificity (31), suggesting non-specific effects may negate changes in task performance. Antidepressants with a significant SSRI component appear to drive bidirectional effects between the two tasks.

### Chronic treatment induces distinct behavioural effects to acute treatment

While work has demonstrated changes in brain region activity relevant to apathy in healthy participants after one week of citalopram treatment (4) it is less clear when behavioural symptoms emerge. In direct contrast to its acute effects, chronic escitalopram increased nesting material shuttled in the EBF task, regardless of effort contingency in the effort curve paradigm. This increase in foraging behaviour is consistent with an increase in motivational state and highlights how treatment length can produce distinct behavioural effects reflecting neurobiological changes associated with longer-term elevated serotonergic exposure. In contrast, trials and chow consumed were decreased in the EfR task, which may be driven by an overall reduction in appetite rather than a specific motivational deficit. This is consistent with reports that escitalopram use is associated with decreased appetite (32). Appetite was not specifically tested, however this highlights limitations associated with interpretation of food-motivated tasks. Chronic venlafaxine showed no effect on motivation in the effort curve paradigm or the EfR task. This difference in behavioural profile compared to chronic escitalopram may be due to differences in serotonergic/noradrenergic balance induced by drug treatment. This balance may have relevance to apathy behaviour as SNRIs are associated with a lower prevalence of apathy than SSRIs (33).

The set up of the EBF task generates some conflict in requiring the mouse to leave a safe and enclosed area to forage and thus may provide additional insight into affective reactivity changes and its impacts on foraging behaviour. A larger (and thus more aversive) area reduced total and % nesting material shuttled in both groups indicating an overall suppression of foraging behaviour in response to an aversive environment. However, previous work has shown that change in % material shuttled to the home area in the aversive context may be sensitive to modulation of affective reactivity in phenotypic models such as healthy ageing and chronic corticosterone treatment (7). A blunting in affective reactivity could mitigate the impact on foraging in an aversive environment. There was a trend towards chronic venlafaxine treated animals shuttling more nesting material in an aversive environment suggesting it may have an emotional blunting effect. A greater range of affective tests is necessary to probe this further.

Contrasting our original hypothesis, elevation in motivation and lack of emotional blunting observed with chronic escitalopram is inconsistent with apathy behaviour. SSRI-induced apathy syndrome seems to be dose and duration dependent as it becomes reversible with lowering the dosage or discontinuation of the drug (33). Therefore, our dose and selected time window may not have been sufficient to induce apathy-relevant behaviours in healthy mice. However, EBF findings do reveal that dose and time course-induced differences in serotonergic modulation may have a complex relationship with apathy behaviours that should be further investigated. The EfR task appeared to have limited sensitivity to chronically-induced behavioural changes which may relate to mechanistic differences between tasks as described above. It is critical that future work investigate the underlying neurobiological differences between these two tasks including regional differences to better understand divergence in drug effect and its relationship with different constructs of motivated behaviour.

## Conclusion

Using a range of antidepressants we show that acute treatment has opposing behavioural effects depending on choice of motivation-based task. While antidepressant treatment impaired aspects of effortful foraging behaviour, it improved effortful conditioned responding. In contrast, chronic treatment with escitalopram increased effortful foraging behaviour but induced a general impairment in the EfR task. Chronic venlafaxine treatment had limited effect on task output. These findings reveal two important points; the first is that antidepressant effect on motivation and other apathy-related behaviours may be dose and time-course dependent which may have important clinical implications. Secondly, two tasks of effort-based motivation (innate versus conditioned) produce very different outcomes which has critical translational importance relating to interpretation of drug effect. Ultimately, this work highlights the need to understand how these types of motivation differ, and how they relate to the behavioural repertoire that makes up our experience of motivation in daily life.

## Data statement

Data will be made available on the Open Science Framework upon acceptance.

## Acknowledgements

The authors would like to gratefully acknowledge Julia Bartlett for their technical assistance in behavioural testing, and the University of Bristol Animal Services Unit for care of the mice.

## Contributions

MGJ – Conception and design of work, analysis and interpretation of data, writing and editing manuscript, final approval of draft. FX – acquisition of data, analysis and interpretation of data, writing and editing manuscript, final approval of draft. CM – acquisition of data, analysis and interpretation of data, writing and editing manuscript, final approval of draft. LL-acquisition of data, final approval of draft. ESJR-writing and editing manuscript, final approval of draft.

