## Supplementary materials for "Conditioned versus innate effort-based tasks reveal divergence in antidepressant effect on motivational state in male mice"

**S1. Cohort summary**

| Cohort | Acute v Chronic | Task | Drug | Age (weeks) |
| --- | --- | --- | --- | --- |
| CM1 (n=16) | Acute | EfR | Escitalopram | 16 |
| | | | Citalopram | 18 |
| | | | Fluoxetine | 22 |
| | | | Venlafaxine | 24 |
| | | | Reboxetine | 26 |
| | | | Sertraline | 28 |
| | | | Vortioxetine | 30 |
| MJ17 (n=16) | Acute | EBF | Escitalopram | 23 |
| | | | Citalopram | 25 |
| | | | Fluoxetine | 27 |
| | | | Venlafaxine | 29 |
| | | | Reboxetine | 32 |
| | | | Sertraline | 34 |
| | | | Vortioxetine | 36 |
| FX1 (n=24) | Chronic | Treatment start | Escitalopram | 15 |
| | | EfR | | 17 |
| | | EBF – 4hr | | 18 |
| | | EBF – Effort curve | | 19-20 |
| | | EBF – Affective reactivity | | 21 |
| CM2 (n=24) | Chronic | Treatment start | Venlafaxine | 14 |
| | | EfR | | 16 |
| | | EBF – 4hr | | 17 |
| | | EBF – Effort curve | | 18-19 |
| | | EBF – Affective reactivity | | 19-20 |

**S1**. *Summary of n numbers and ages per experiment* EBF=effort based foraging task, EfR=effort for reward task, NSFT= novelty supressed feeding task

**S2. Additional husbandry information**

Cages were enriched with a small red house, a tube, a tube suspended from the ceiling, a wooden chew, nesting material and a nestlet. Mice didn't start training until one week after their arrival and were habituated to cup-handling before training began.

**S3. Effort for Reward training stages**

In all stages mice underwent one 30-min session per day. In the first stage, mice learned to associate the magazine with the automatic delivery of a reward over two sessions. Mice then progressed to six sessions of continuous reinforcement training (CRF). Here, a response could be made in either of the two nose poke apertures, resulting in the delivery of a reward pellet. The mice then progressed to fixed ratio (FR) training where a response could be made in either the left or right nose poke aperture only, counterbalanced across the cohort.

**S4. Effort for Reward secondary task outputs**

| Measure | Description |
|---|---|
| Number of trials | The number of completed high-effort trials which resulted in high-reward delivery. |
| Chow consumed | The amount of chow eaten from the bowl within the session (grams) |
| *Chow bouts* | *The number of times the mouse visits the bowl throughout the session.* |
| *Duration at chow bowl* | *The average duration of the bouts at the chow bowl (s).* |
| *Latency to chow bowl* | *The time taken for the mouse to interact with the chow bowl for the first time within the session (s).* |
| *Latency to complete first trial* | *Time taken for the mouse to complete the first trial to receive a reward pellet (s).* |

***Table 1 Output measures of the effort for reward task.*** *Italicised entries refer to output measures relating to chow bowl from video analysis*

**S5. Acute and chronic experimental designs for the Effort for Reward task**

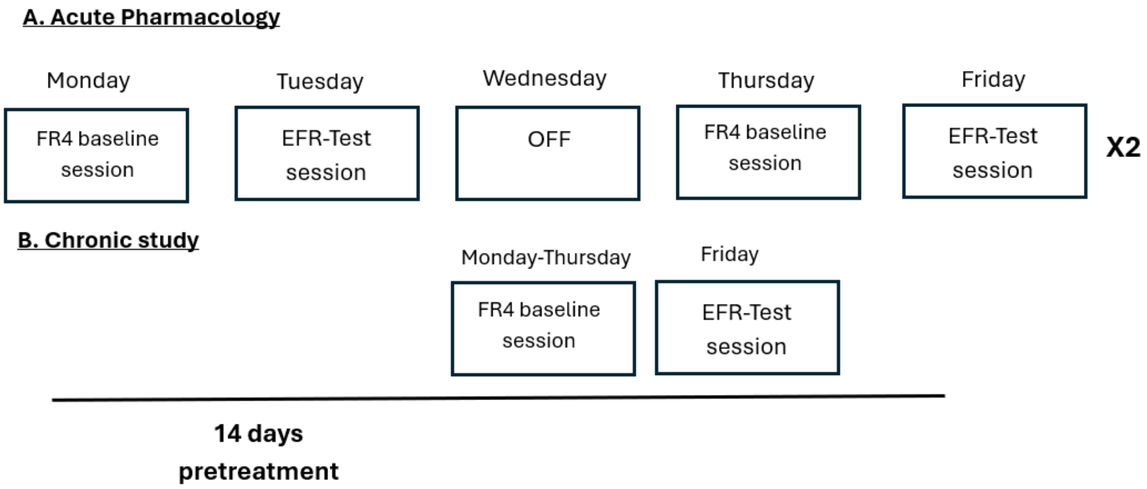


***S5**: **Experimental design of the EFR task acute and chronic pharmacological studies**. **A.*** *Each mouse received every dose of the drug in a within-subject, counterbalanced design spread across 4 test sessions (one dose per session). Each test session was preceded by an FR4 baseline day and a day off on the Wednesday.* ***B.*** *Mice received 14 days of treatment before undergoing 4 days of re-baselining and a subsequent Effort for Reward test session.*

**S6. Drug vehicles**

| Drug | Dose Range | Mouse AED | Pretreatment time | Administration Route | Vehicle |
|---|---|---|---|---|---|
| **Escitalopram** | 1-10 mg/kg | 1.59-3.18 mg/kg | 1h | Oral | 20% condensed milk |
| **Citalopram** | 1-10 mg/kg | 3.18mg/kg | 1h | Oral | 20% condensed milk |
| **Fluoxetine** | 1-10 mg/kg | 3.18mg/kg | 4h | Oral | 50% condensed milk |
| **Venlafaxine** | 1-10 mg/kg | 5.95mg/kg | 1h | Oral | 20% condensed milk |
| **Reboxetine** | 0.3-3 mg/kg | 1.27mg/kg | 30 mins | Oral | 20% condensed milk |
| **Sertraline** | 1-10 mg/kg | 3.97-7.94 mg/kg | 4h | Oral | 50% condensed milk |
| **Vortioxetine** | 0.3-3 mg/kg | 0.8-1.59 mg/kg | 2hours | Oral | 50% condensed milk |

***Table 2: Summary of acute drug studies, listed in the order of administration*** *AED=Animal Equivalent Dose*

**S7. Habituation details**

**First session:** The mouse was placed in the home area of the forage arena for 10 minutes. The forage arena contained woodchip only. The mouse was able to freely explore the entirety of the arena during this time. Woodchip was exchanged between mice. The arena was cleaned with water and ethanol once all mice had run the session.

**Second and third session:** This was repeated on the second and third day for a period of 5 minutes only.

**S8A. Arena details**

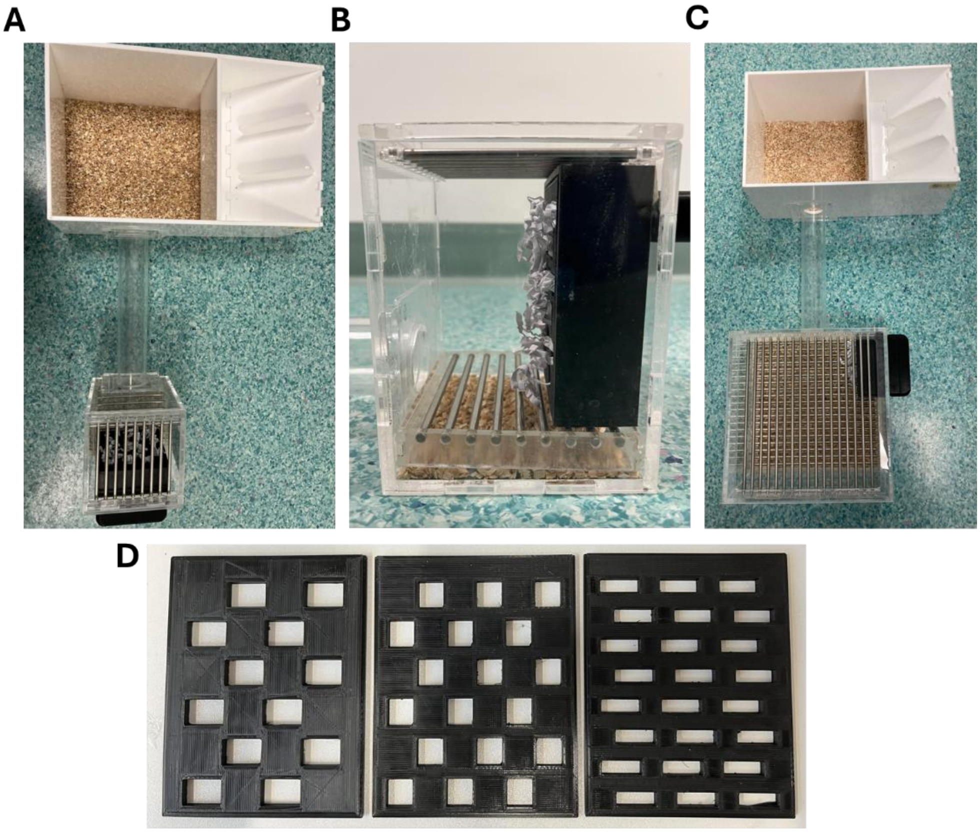

***S8A. Effort-Based Forage task arenas. A** Shows the standard arena set up containing a white Perspex home area, with a tube connecting to a forage area where the nesting box is fixed with a magnet bar. The forage area is made from clear Perspex and has a steel barred lid to expose the mouse to standard unit airflow, creating more 'open' conditions. The floor is also barred and lined with woodchip to catch fallen faecal pellets and promote shuttling of nesting material to the home area. **B** Shows a side view of the nesting material box. 18g Sizzlenest is accessible in the nesting box via apertures in the face plate. During testing the mice can freely choose to traverse the tube to forage nesting material and bring it back to the home area. **C** Shows the enlarged forage arena set up, used to test affective reactivity by providing a more open and novel environment in which to forage from the nesting box. The nesting box is positioned equidistant to the tube opening to that of the standard arena to prevent any confounding impact of extra distance required to shuttle nesting material. **D** Shows the different aperture sizes in the nesting box faceplates used to generate the effort curve. The widest apertures (left) are used during the habituation session. The moderate apertures (middle) are used for drug testing. Figure is adapted from Xeni et al 2024.*

**S8B. Arena details**

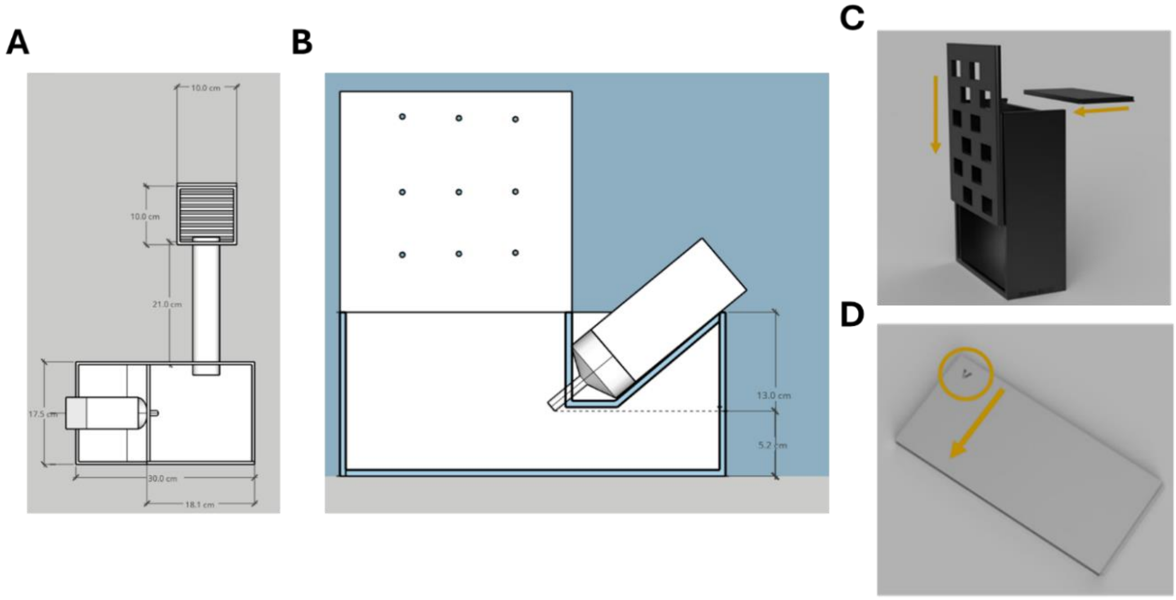


***S8B. Dimensional renderings of the Effort-Based Forage arena and nesting material box. A*** *A top down view of the standard arena set up with associated dimensions.* ***B*** *A side view of the home area showing water bottle holder placement and associated dimensions. Note-lid is not hinged.* ***C*** *The nesting box is printed as separate components (main body, face plate, lid) which slot together as shown above.* ***D*** *The nesting box lid has a small printed arrow showing the direction it should be slotted into the nesting box. Figure adapted from Xeni et al 2024.*

**S8C. Arena details**

| Component | Dimensions |
|---|---|
| **Home area** | W 17.5 cm, L 30.0 cm, H 13.0 cm |
| **Home area lid** | W 18.1 cm, L 18.1 cm, H 18.1 cm |
| **Connecting tube** | L 21.0 cm |
| **Forage area** | W 10.0 cm, L 10.0 cm, H 13.0 cm |
| **Forage area (large)** | W 21.0 cm, L 21.0 cm, H 14.0 cm |
| **Nesting material box body (3D printed)** | W 3.5 cm, L 8.0 cm, H 10.0 cm |
| **Nesting material box face plate (3D printed)** | W 0.3 cm, L 7.8 cm, H 10.0 cm |
| **Nesting material box lid (3D printed)** | W 3.5 cm, L 7.8 cm, H 0.3 cm |
| **Magnet bar (3D printed)** | W 2.0 cm, L 8.0cm, H 3.0 cm |
| **'Easy apertures' (1.5 cm$^2$)** | W 1.5 cm H 1 cm |
| **'Moderate apertures' (1 cm$^2$)** | W 1 cm H 1 cm |
| **'Difficult apertures' (0.75 cm$^2$)** | W 1.5 cm H 0.5 cm |

**S8: *Dimensions of Effort-Based Forage arena components.*** *Note 3D print files are available via Xeni et al 2024.*

**S9. Effort-based Forage task outputs**

| **Measure** | **Description** |
|---|---|
| Total nesting material shuttled to the home area (g) | The amount of nesting material pulled from the nesting box (foraged) throughout the session and taken through the tube to the home area (grams) |
| % nesting material foraged taken to the home area | Of the total bedding foraged, how much is transported to the home area instead of left on the floor of the forage area, expressed as percentage |
| *AUC (Activity x time bout)* | *The general activity of the mice during the task within the forage area. A percentage of movement within 10 secs was calculated and outputted in 10 second time bins. Prism uses the trapezoidal method to calculate the area-under-the-curve. Each connecting segment's area defines a trapezoid. Prism uses the area of the comparable rectangle to determine the area of each trapezoid. The total area of all the rectangles equals the area under the curve.* |

*S9 **Output measures of the effort-based forage task.** Italicised entries refer to output measures relating to activity within the forage area from sensor analysis. AUC= area under the curve.*

**S10. Acute and chronic experimental designs for the Effort-Based Forage task**

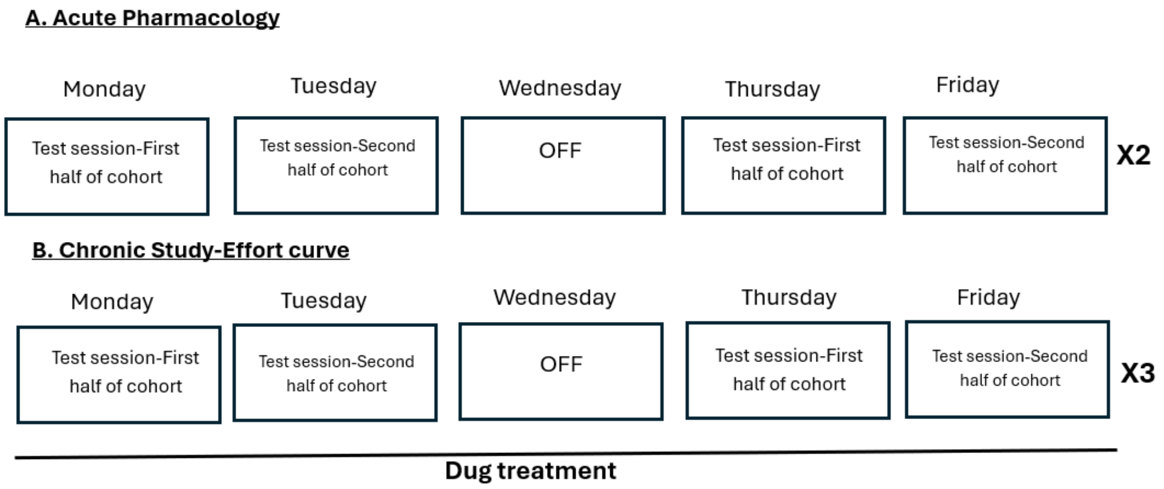


***S10: Experimental design of effort-based foraging task acute and chronic pharmacological design. A.** Each mouse received each dose and appropriate vehicle in a within-subject counterbalanced design, over four test sessions. **B.** Each mouse encountered the 3 different sizes of the aperture in a fully randomised, within-subject design over six test sessions (3 sizes x 2 days per session) with a final sample size (n) of 24 for each size.*

**S11. General activity monitoring**

As previously described by (1), the system was built in-house by MGJ to monitor home cage activity in individually-housed mice, based on a system previously developed by (2). 4 activity sensors with built-in amplifiers (AMN 2,3,4 series Motion Sensor, Panasonic) were read using an Arduino Mega 2560 with an Arduino Mega daughter board (SchmartBoard, Mouser). Data was read onto an SD card as CSV file using an SD card adaptor (TFT LCD w/microSD Breakout, Adafruit) and exact time was outputted alongside the data using a RealTime clock (Clock & Timer Development Tools PCF8523 RTC for RPi, Adafruit). Each sensor was placed above the middle of the forage area, approximately 5 cm from the top of the cage. Movement was detected by the sensor every 100 msec. A percentage of movement within 10 secs was calculated and outputted in 10 second time bins. Area under the curve was calculated using time bin x activity count. Code to run the sensor system was adapted from (2) and can be found here https://github.com/meganjackson13/Sensor-system-code.

**S12. Experimental design for home cage activity testing**

| Stage | Description |
|---|---|
| **Habituation** | Mice habituated to the recording room for 30 mins. |
| **Dosing** | Mice treated with the drug and remained in the recording room for 1 hour more, without being recorded to match pre-treatment time in the main study. |

| | |
|---|---|
| **Test** | Sensor system recorded mice activity for 2 hours. |

**S12:** ***Training protocol of home cage activity monitoring.***

**S13. Statistical exclusions**

| Experiment | Measure | Statistical/technical exclusion in dataset |
|---|---|---|
| **Escitalopram** | | |
| ***EBF task*** | Total taken through | N=1 vehicle, n=1 10mg/kg outlier |
| | %total taken to main box | N=1 vehicle, n=1 1mg/kg outlier, n=1 3mg/kg outlier |
| | General locomotor activity | N=1 vehicle, n=1 3mg/kg outliers |
| ***EfR task*** | Number of trials | No exclusions |
| | Chow consumed | N=1 3mg/kg outlier, n = 1 vehicle outlier |
| | Number of chow bouts | N=1 vehicle outlier, n=1 vehicle technical failure, n=1 1mg/kg technical failure, n=2 3mg/kg outliers, n=1 10mg/kg outlier |
| | Average duration | N=1 vehicle technical failure, n=1 1mg/kg technical failure, n=1 1mg/kg outlier, n=1 10mg/kg outlier |
| | Latency to chow | N=1 vehicle pre-excluded, n=1 vehicle outlier, n=1 1mg/kg pre-excluded, n=1 1mg/kg, n=1 3mg/kg, n=1 10mg/kg outliers |
| | Latency to pellet | N=1 vehicle pre-excluded, n=1 vehicle outlier, n=1 1mg/kg pre-excluded, n=1 1mg/kg, n=1 3mg/kg, n=2 10mg/kg outliers |
| **Citalopram** | | |
| ***EBF task*** | Total taken through | N=1 1mg/kg, n=1 3mg/kg, n=1 10mg/kg outliers |
| | %total taken through | N=1 vehicle, n=1 1mg/kg, n=1 3mg/kg outliers |
| | General locomotor activity | N=1 1mg/kg outlier |
| ***EfR task*** | Number of trials | No exclusions |
| | Chow consumed | N=1 1mg/kg, n=1 10mg/kg outliers |

| | | |
|---|---|---|
| | Number of chow bouts | N=1 vehicle outlier |
| | Average duration | N=1 vehicle, n=1 1mg/kg, n=1 10mg/kg outliers |
| | Latency to chow | N=1 vehicle, n=1 1mg/kg, n=1 3mg/kg, n=1 10mg/kg outliers |
| | Latency to pellet | N=1 1mg/kg, n=1 3mg/kg, n=1 10mg/kg outliers |
| **Sertraline** *EBF task* | Total taken through | N=1 1mg/kg outlier, n=1 3mg/kg |
| | %taken through | N=1 vehicle, n=1 1mg/kg, n=1 3mg/kg, n=1 10mg/kg outliers |
| | General locomotor activity | N=1 full exclusion due to 4 outliers, n=1 3mg/kg, n=1 10mg/kg outliers |
| *EfR task* | Number of trials | N=1 10mg/kg outlier |
| | Chow consumed | No exclusions |
| | Number of chow bouts | N=1 vehicle, n=1 1mg/kg, n=1 3mg/kg outliers |
| | Average duration | N=1 1mg/kg, n=1 3mg/kg, n=1 10mg/kg outliers |
| | Latency to chow | N=1 vehicle, n=1 1mg/kg, n=2 3mg/kg, n=1 10mg/kg outliers |
| | Latency to pellet | N=1 full exclusion due to 3 outliers, n=1 3mg/kg outlier |
| **Fluoxetine** | Total taken through | N=1 10mg/kg outlier |
| *EBF task* | %taken through | N=2 vehicle, n=1 1mg/kg outlier, n=1 3mg/kg outlier |
| | General locomotor activity | N=1 3mg/kg, n=1 10mg/kg outliers |
| *EfR task* | Number of trials | N=1 1mg/kg outlier, n=1 3mg/kg outlier |
| | Chow consumed | N=1 vehicle |
| | Number of chow bouts | No exclusions |
| | Average duration | N=1 vehicle, n=2 1mg/kg, n=1 10mg/kg outliers |
| | Latency to chow | N=2 vehicle, n=1 1mg/kg, n=2 3mg/kg, n=1 10mg/kg outliers |
| | Latency to pellet | N=1 full exclusion due to three outliers |

| | | |
|---|---|---|
| **Venlafaxine**<br><br>***EBF task*** | Total taken through | N=1 vehicle, n=1 1mg/kg outlier |
| | %taken through | N=1 vehicle, n=1 1mg/kg, n=1 3mg/kg, n=1 10mg/kg outliers |
| | General locomotor activity | Technical failure |
| ***EfR task*** | Number of trials | N=1 (failed due to technical error) vehicle, n=1 3mg/kg outlier, n=1 10mg/kg outlier |
| | Chow consumed | N=2 (1 failed due to technical error) vehicle, n=2 1mg/kg, n=1 3mg/kg outliers |
| | Number of chow bouts | N=1 (failed due to technical error) vehicle |
| | Average duration | N=1 (failed due to technical error) vehicle, n=1 excluded |
| | Latency to chow | N=1 (failed due to technical error) vehicle, n=1 vehicle, n=1 1mg/kg, n=1 3mg/kg, n=1 10mg/kg outliers |
| | Latency to pellet | N=1 (failed due to technical error) vehicle, n=1 vehicle, n=1 1mg/kg, n=1 3mg/kg, n=2 10mg/kg outliers |
| **Reboxetine**<br><br>***EBF taks*** | Total taken through | N=1 vehicle, n=1 3mg/kg outlier |
| | %taken through | N=2 vehicle, n=1 0.3mg/k, n=1 1mg/kg, n=1 3mg/kg outliers |
| | General locomotor activity | N=1 3mg/kg outlier |
| ***EfR task*** | Number of trials | No exclusions |
| | Chow consumed | N=1 3mg/kg |
| | Number of chow bouts | N=1 vehicle, n=1 3mg/kg outliers |
| | Average duration | N=1 vehicle, n=1 0.3mg/kg, n=1 1mg/kg, n=1 3mg/kg outliers |

| | | |
|---|---|---|
| | Latency to chow | N=1 full exclusion due to 4 outliers, n=1 1mg/kg outlier |
| | Latency to pellet | N=1 vehicle, n=1 0.3mg/kg, n=1 1mg/kg outliers |
| **Vortioxetine** | Total taken through | N=2 3mg/kg outliers |
| ***EBF task*** | %taken through | N=1 vehicle, n=1 0.3mg/kg, n=2 1mg/kg, n=2 3mg/kg outliers |
| | General locomotor activity | N=1 vehicle, n=1 1mg/kg, n=1 3mg/kg outliers |
| ***EfR task*** | Number of trials | N=1 0.3mg/kg outlier, n=1 1mg/kg outlier |
| | Chow consumed | N=1 vehicle, n=1 3mg/kg outlier |
| | Number of chow bouts | N=1 vehicle outlier |
| | Average duration | N=1 1mg/kg, n=1 3mg/kg outliers |
| | Latency to chow | N=1 vehicle, n=2 0.3mg/kg, n=1 1mg/kg, n=1 3mg/kg outliers |
| | Latency to pellets | N=1 full exclusion due to 3 outliers |
| **Chronic ESC** ***EfR task*** | Number of trials | N=1 (+) outlier |
| | Chow consumed | No exclusions |
| | Number of chow bouts | No exclusions |
| | Average duration | N=1 (+) outlier |
| | Latency to chow | N=1 (+) outlier |
| | Latency to pellet | N=1 (-) outlier |
| ***EBF task: First session*** | Total taken through | N=1(+), n=1 (-) outliers |
| | %taken through | N=1(+), n=1 (-) outliers |
| ***Effort curve*** | Total taken through | N=1 (+) pre-excluded due to box error, n=1 0.75cm (-) outlier, n=1 (+) excluded due to 3 outliers, n=1 0.75cm (+) outlier |

| | | |
|---|---|---|
| | %taken through | N=1 (+) pre-excluded due to box error, n=1 (+) excluded due to 2 outliers, n=1 1.5cm (+) outlier, n=1 0.75cm (-), n=1 1cm (-), n=1 1.5cm (-) outliers |
| ***Big vs std*** | Total taken through | N=1 big (-), n=1 big (+) outliers, n=1 (+) pre-excluded due to box error |
| | %taken through | N=1 (+) pre-excluded due to box error, n=1 (+) full exclusion due to two outliers |
| **Chronic Venlafaxine**<br>***EfR task*** | Number of trials | N=1 (-) (failed due to technical error), n=1 (-), n=1 (+) outliers |
| | Chow consumed | N=1 (-) (failed due to technical error), n=1 (-) outlier, n=1 (+) outlier |
| | Number of chow bouts | N=1 (-) failed due to technical error |
| | Average duration | N=1 (-) failed due to technical error, n=1 (+) outlier |
| | Latency to chow | N=1 (-) (failed due to technical error), n=1 (-), n=1 (+) outliers |
| | Latency to pellets | N=1 (-) failed due to technical errors, n=1 (+) outlier |
| ***EBF task:***<br>***First session*** | Total taken through | N=1 (-), n=1 (+) outliers |
| | % taken through | N=2 (-), n=1 (+) outliers |
| ***Effort curve*** | Total taken through | N=1 pre-excluded due to lack of task engagement, n=1 (+) 1.5cm outlier |
| | % taken through | N=1 pre-excluded due to lack of task engagement, n=1 (-) 0.75cm, n=1 (-) 1cm, n=1 (-) 1.5cm, n=1 (+) 1cm, n=1 (+) 0.75cm outliers |
| ***Big vs std*** | Total taken through | N=1 pre-excluded due to lack of task engagement, n=1 (-) big, n=1 (-) std, n=1 (+) big, n=1 (+) std outliers |
| | % taken through | N=1 pre-excluded due to lack of task engagement, n=1 (-) std, n=1 (+) std outliers |

**S13:** ***Summary of data point exclusions/replacements per experiment and output measure.***

**S14. Acute escitalopram reduced general locomotor activity in the forage area, whereas no such effect was observed with the other SSRIs and non-SSRIs drugs**

Mice were dosed with a range of SSRIs and underwent the EBF task. Escitalopram reduced general locomotor activity in the forage area ($F_{(2.542, 38.14)} = 8.895$, $p = 0.0003$, (3mg/kg, $p = 0.0057$), (10mg/kg, $p = 0.0064$)) (**Fig. S14A**). However, there was no effect of escitalopram on home cage activity (**Fig. S15**). Sertraline and fluoxetine had no effect on general locomotor activity (**Fig. S14B-C**).

Mice were dosed with a range of non-SSRI antidepressants and underwent the EBF task. Reboxetine and vortioxetine had no effect on general locomotor activity in the forage area (**Fig. S14D, E**).

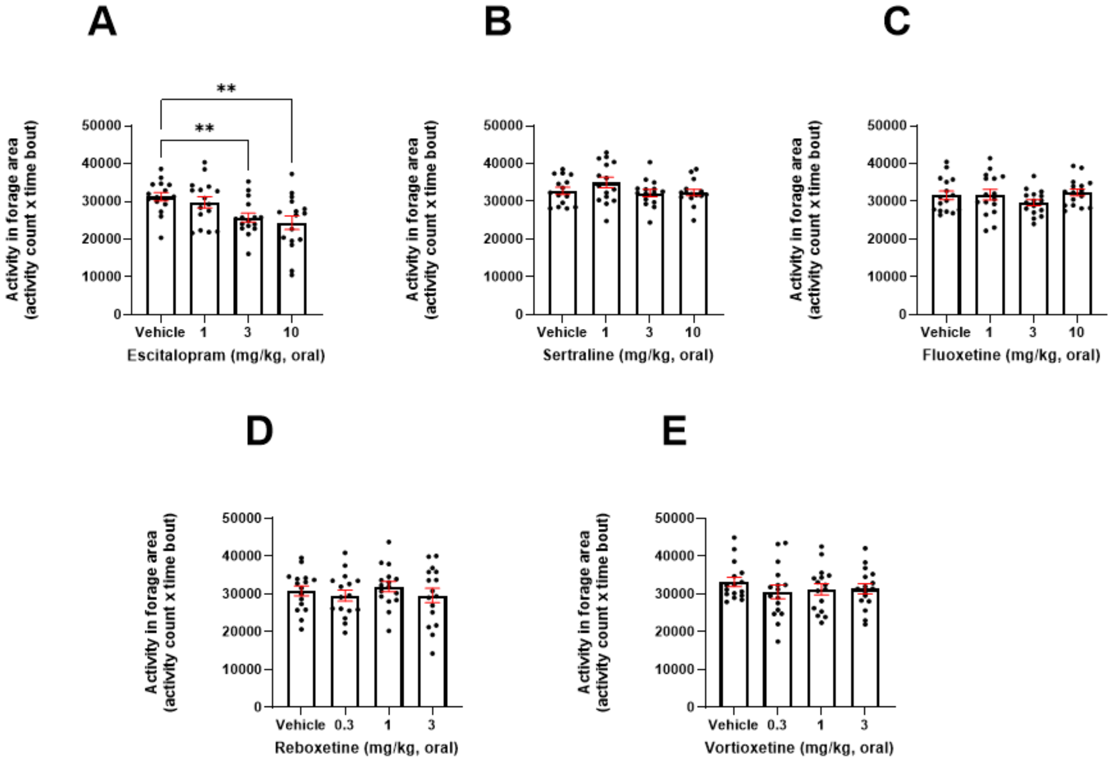


**S14:** ***All drugs with the exception of escitalopram had no effect on general activity in the forage area.*** *Activity was recorded in the forage area of the arena using a passive infrared sensor system.* ***A.*** *3 and 10mg/kg escitalopram reduced general locomotor activity in the forage area ($p < 0.01$).* ***B-F.*** *There was no effect of sertraline, fluoxetine, venlafaxine, reboxetine or vortioxetine on general locomotor activity in the forage area ($p > 0.05$). Error bars are mean ±SEM, one-way ANOVA was performed to detect the drug effect, *$p<0.05$.*

**S15. Home cage activity analysis-escitalopram**

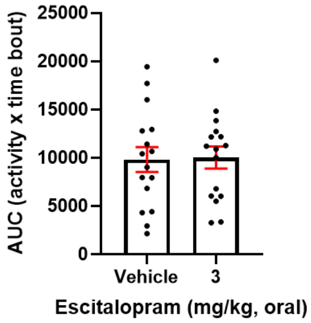


***S15: Escitalopram had no effect in the activity of the mice under home-cage conditions.*** *3mg/kg escitalopram had no effect in the home-cage activity relative to vehicle (p>0.05). Error bars are mean ±SEM, independent samples t-test was performed.*

**S16. Acute escitalopram slightly impairs chow-related behaviour in the EfR task, with no effects observed for other SSRIs**

Mice were dosed with a range of SSRIs and underwent the EfR task. Escitalopram reduced chow bowl bouts ($F_{(2.544, 38.15)}$ = 10.46, p < 0.0001, (10mg/kg, p = 0.0013)) (**Fig. 16A**). Escitalopram trended towards a decrease on the average time spent at the chow bowl ($F_{(2.019, 30.28)}$ = 2.882, p = 0.0711) (**Fig. 16B**). Escitalopram had no effect on latency to first time eating chow (**Fig. 16C)** or latency to complete first trial (**Fig. 16D**).

Sertraline had no effect on any of the secondary bowl output measures (**Fig. 16E-H**).

Fluoxetine had no effect on chow bowl bouts (**Fig. 16I**). Fluoxetine trended towards a decrease on average time spent at the chow bowl ($F_{(2.169, 32.53)}$ = 2.931, p = 0.0636) (**Fig. 16J**), with no effect on the other secondary parameters (**Fig. 16K, 16L**).

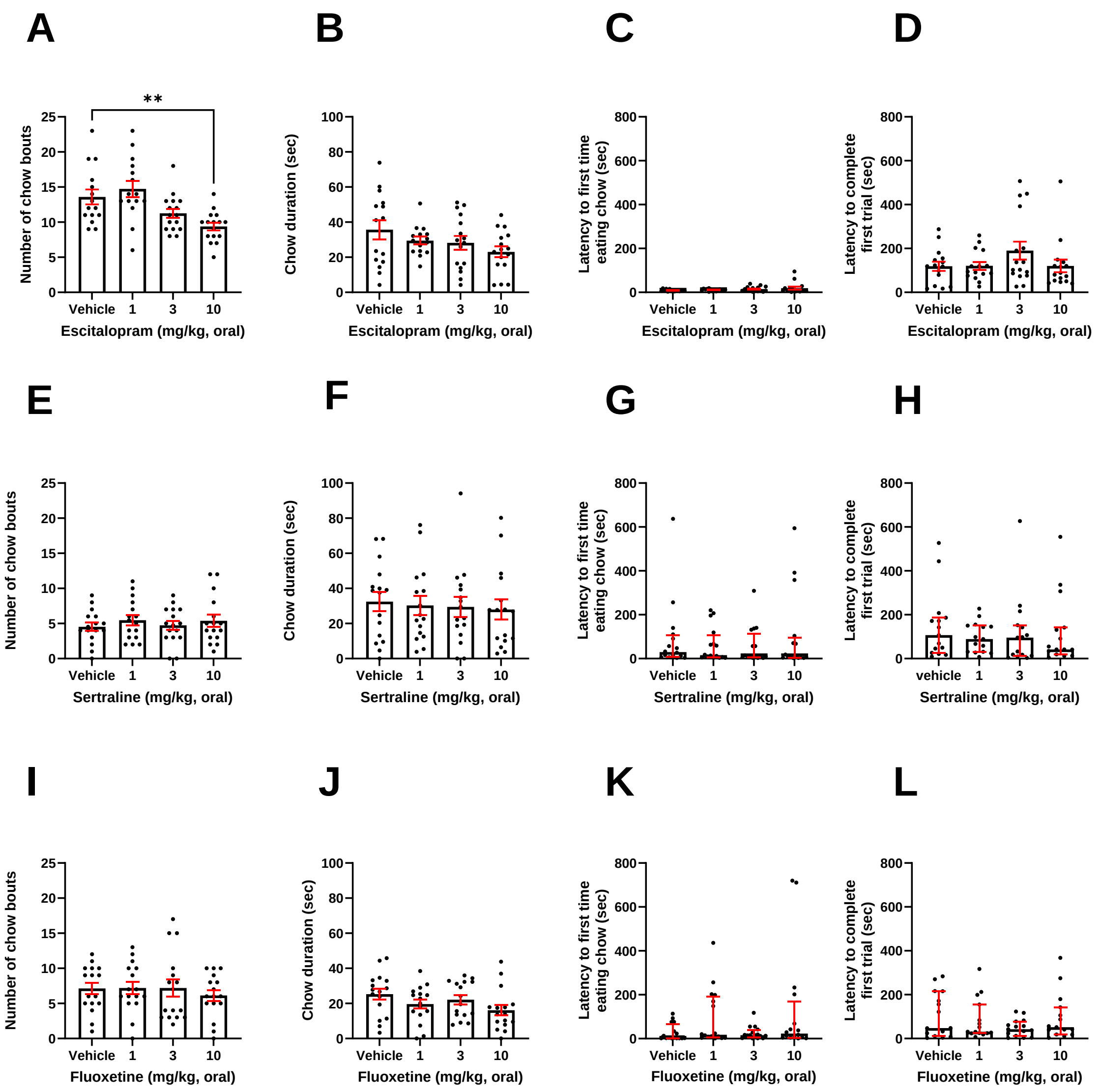


**S16*: Acute escitalopram slightly impairs chow-related behaviour in the EfR task, with no effects observed for other SSRIs.*** *Mice were dosed with a range of SSRIs and underwent the EfR task.* ***A.*** *10mg/kg escitalopram reduced number of chow bouts ($p < 0.01$). Escitalopram did not affect* ***B.*** *average time spent at chow bowl ($p > 0.05$),* ***C.*** *latency to first time eating chow ($p > 0.05$) or* ***D.*** *latency to complete first trial ($p > 0.05$). Sertraline did not affect* ***E.*** *number of chow bouts ($p > 0.05$),* ***F.*** *average time spent at chow bowl ($p > 0.05$),* ***G.*** *latency to first time eating chow ($p > 0.05$) or* ***H.*** *latency to complete first trial ($p > 0.05$). Fluoxetine did not affect* ***I.*** *number of chow bouts ($p > 0.05$),* ***J.*** *average time spent at chow bowl ($p > 0.05$),* ***K.*** *latency to first time eating chow ($p > 0.05$) or* ***L.*** *latency to complete first trial ($p > 0.05$). Error bars are mean ±SEM, one-way ANOVA was performed to detect the drug effect, **$p<0.01$.*

**S17. Acute citalopram treatment exerts opposing effects in the EfR and EBF task**

Mice were dosed with citalopram and underwent the EBF and EfR task. Citalopram reduced nesting material shuttled ($F_{(2.424,\ 36.36)}$ = 35.01, p<0.0001, (3mg/kg, p= 0.0179), (10mg/kg, p<0.0001)) (**Fig.17A**). Citalopram also reduced % shuttled (*Q = 11.40,* p=0.0098, (10mg/kg, p=0.0030) (**Fig.17B)** Forage area activity was not affected **(Fig. 17C).**

Citalopram increased number of trials completed in the EfR task ($F_{(2.580,\ 38.70)}$ = 38.68, p < 0.0001, (3mg/kg, p=0.0005), (10mg/kg, p < 0.0001)) (**Fig.17D**). Citalopram trended towards a decrease in the amount of chow consumed ($F_{(1.809,\ 27.13)}$ = 2.600, p = 0.0973) (**Fig.17E**). Citalopram had no effect on latency to complete the first trial **(Fig. S17F),** number of bouts at the chow bowl **(Fig. S17G),** or average duration spent at the chow bowl **(Fig. S17H)**. Citalopram had a main effect on latency to first time eating chow (*Q=8.925*, p=0.0303), however post-hoc analysis didn't reach significance **(Fig. S17I)**.

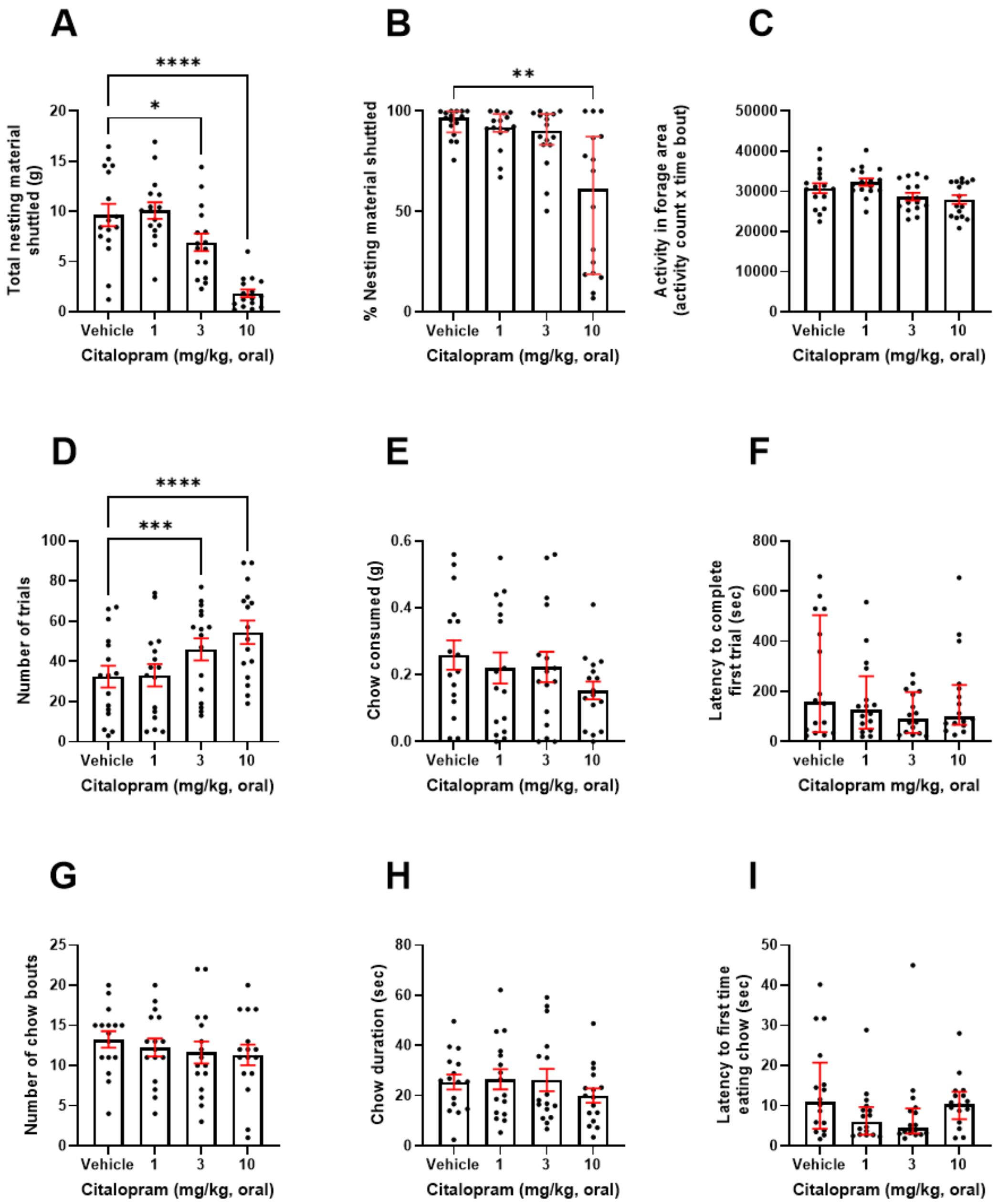


**S17:** ***Acute citalopram treatment exerts opposing effects in the EfR and EBF task.*** *Mice were dosed with 1-10mg/kg citalopram and underwent the EBF or the EfR task.* ***A.*** *3 and 10mg/kg citalopram caused a reduction in the total nesting material shuttled (p<0.05, p<0.0001), B. 10mg/kg citalopram caused a reduction on % of nesting material shuttled (p < 0.01)* ***C.*** *Citalopram had no effect on general locomotor activity within the forage area (p > 0.05)* ***D.*** *3 and 10mg/kg citalopram caused an increase in the number of high-effort, high-value trials performed (p < 0.001, p<0.0001)* ***E.*** *with no effect on latency to complete the first trial (p > 0.05)* ***F.*** *10mg/kg citalopram caused a reduction on the amount of chow consumed (p < 0.05), with no effect on* ***G.*** *the number of chow bouts (p > 0.05),* ***H.*** *the average time spent at the chow bowl (p > 0.05), or* ***I.*** *the latency to approach the chow bowl (p > 0.05). Error bars are mean ±SEM, one-way ANOVA was performed to detect drugs effect, *p<0.05, **p<0.01, ***p<0.001, ****p<0.0001.*

**S18. Acute reboxetine slightly increased chow-related behaviour in the EfR task, with no effects observed for other non-SSRIs**

Mice were dosed with a range of non-SSRIs and underwent the EfR task. Venlafaxine had no effect on chow bowl bouts (**Fig. S18A**), average time spent at chow bowl (**Fig. S18B**), latency to first time eating chow (**Fig. S18C**) or latency to complete the first trial (**Fig. S18D**).

Reboxetine had a main effect on number of bouts at the chow bowl ($F_{(2.550,\ 35.70)}$ = 3.142, p = 0.0442). 3mg/kg reboxetine trended towards decreased number of bouts at the chow bowl (p = 0.0851) (**Fig. S18E**). Reboxetine increased the average time spent at chow bowl ($F_{(2.870,\ 40.18)}$ = 3.563, p = 0.0239, (3mg/kg, p = 0.0219)) (**Fig. S18G**), with no effect on the other measured parameters (**Fig. S18G, H**).

Vortioxetine had no effect on any of the secondary bowl output measures (**Fig. S18I-L**).

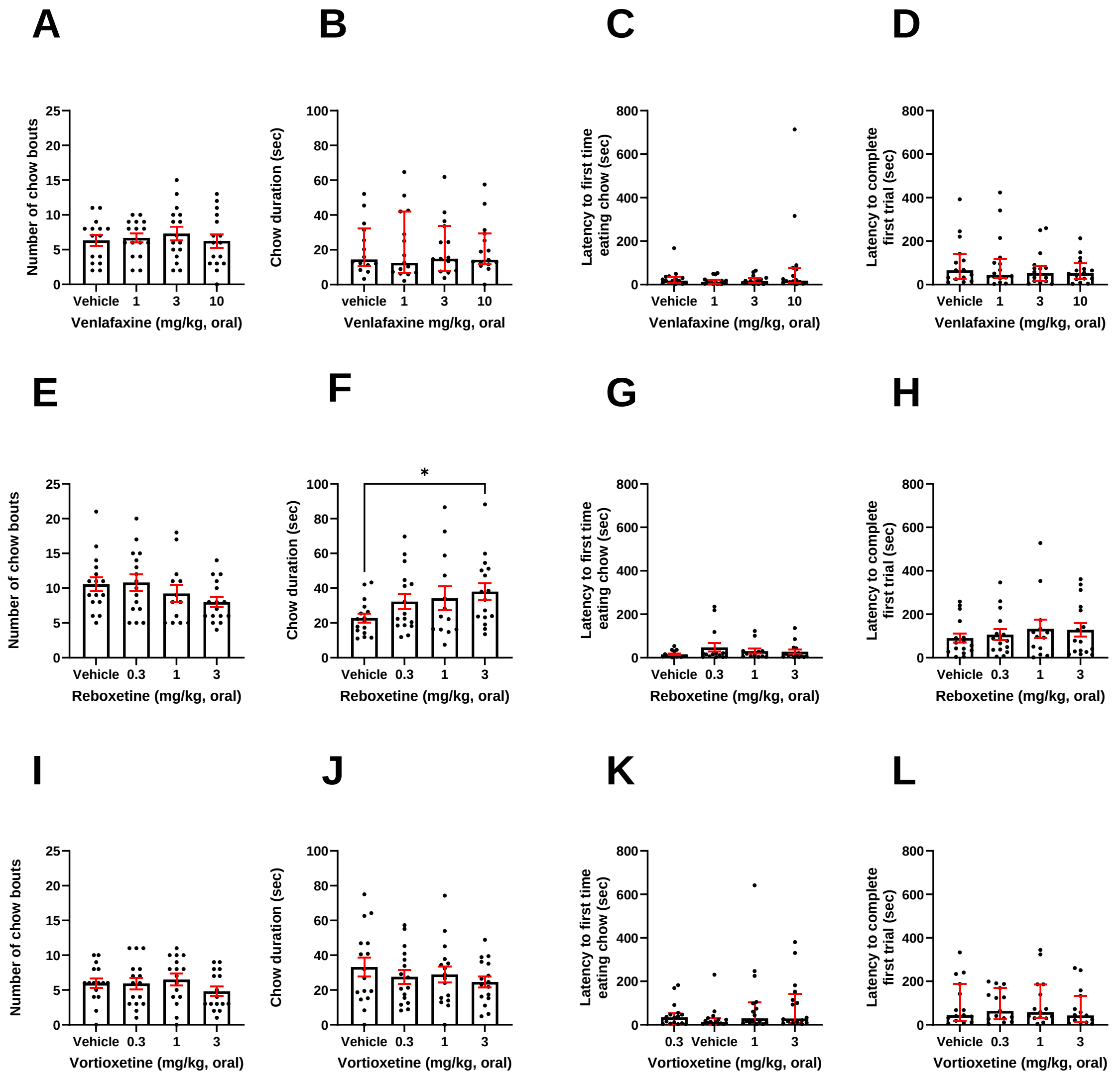

**S18***: **Acute reboxetine slightly increased duration at the chow bowl in the EfR task, with no effects observed for other non-SSRIs.** Mice were dosed with a range of non-SSRIs and underwent the EfR task.* ***A.*** *Venlafaxine did not affect number of chow bouts (p > 0.05)* ***B.*** *average time spent at chow bowl (p > 0.05),* ***C.*** *latency to first time eating chow (p > 0.05) or* ***D.*** *latency to complete first trial (p > 0.05). Reboxetine did not affect* ***E.*** *number of chow bouts (p > 0.05).* ***F.*** *3mg/kg reboxetine increased the average time spent at chow bowl (p < 0.05), with no effect on* ***G.*** *latency to first time eating chow (p > 0.05) or* ***H.*** *latency to complete first trial (p > 0.05). Vortioxetine did not affect* ***I.*** *number of chow bouts (p > 0.05),* ***J.*** *average time spent at chow bowl (p > 0.05),* ***K.*** *latency to first time eating chow (p > 0.05) or* ***L.*** *latency to complete first trial (p > 0.05). Error bars are mean ±SEM, one-way ANOVA was performed to detect the drug effect, *p<0.05.*

**S19. Chronic treatment did not impact initial foraging during habituation**

There was no effect of chronic treatment with either escitalopram or venlafaxine on total nesting material shuttled or % shuttled during the initial 4 hour foraging habituation session (p>0.05) (**Fig.S19A-D**).

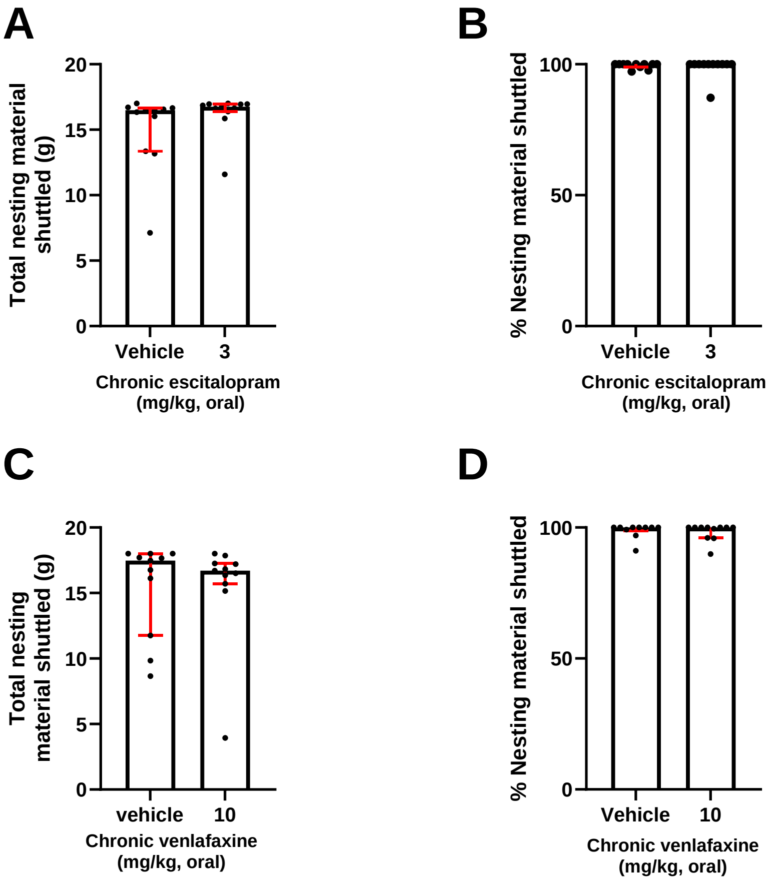


***S19. Chronic treatment with escitalopram or venlafaxine did not impact foraging during habituation. A** Escitalopram did not impact total nesting shuttled or **B** % nesting material shuttled. **C** Venlafaxine did not impact total nesting material foraged or **D** % nesting material shuttled (p>0.05). Bars are median ± interquartile range with data points overlaid.*

**S20. Chronic treatment had limited impact on chow-related behaviour**

Mice were treated with 3mg/kg escitalopram for 2 weeks and underwent the EfR task. Chronic escitalopram had no effect on number of chow bouts (**Fig. S20A**). Treated mice spent less time at the chow bowl, compared to the vehicle ($t_{(21)}$ = 2.259, p = 0.0346) (**Fig. S20B**), with no effect of treatment on latency to first time eating chow (**Fig. S20C**), or latency to complete the first trial (**Fig. S20D**).

Mice were treated with 10mg/kg venlafaxine for 2 weeks and underwent the EfR task. Chronic venlafaxine had no effect on number of chow bouts (Fig. S19E). There was no change in average time spent at the chow bowl ($t_{(20)}$ = 1.665, p = 0.1116) (**Fig. S19F**), and no effect of treatment on latency to first time eating chow (**Fig. S20G**), or latency to complete the first trial (**Fig. S20H**).

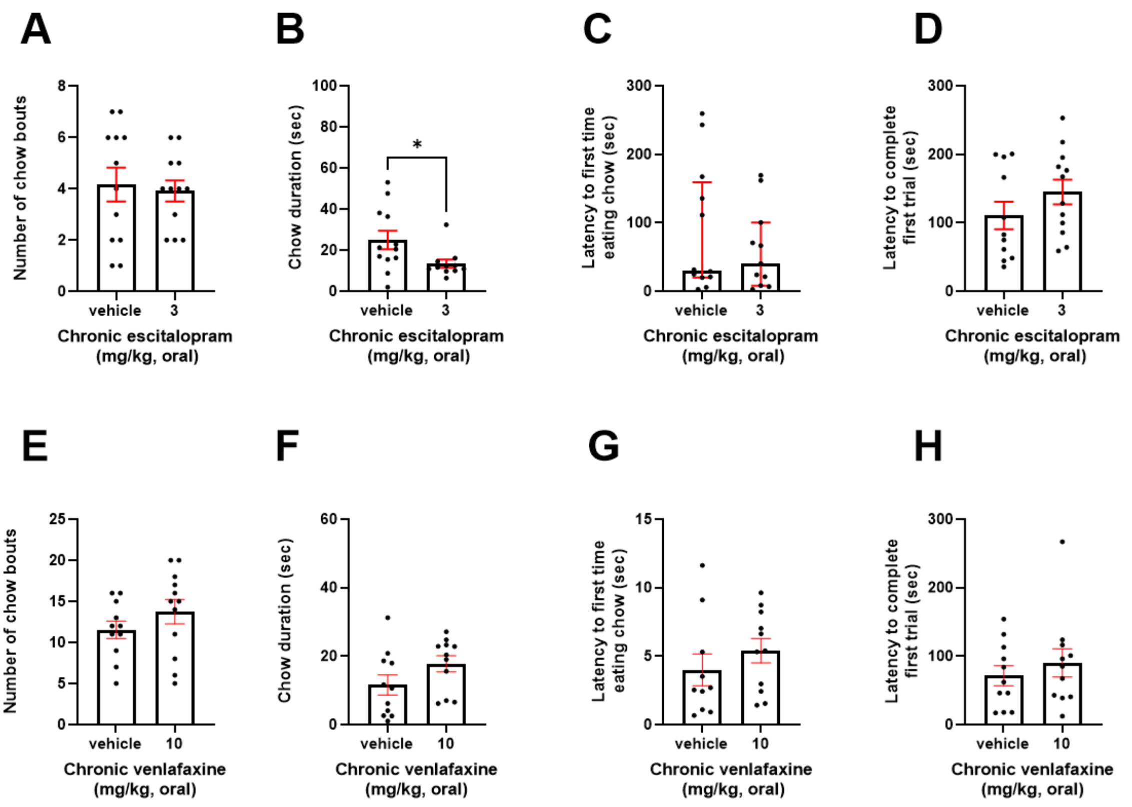


**S20:** ***Chronic escitalopram slightly impaired chow consumed related behaviour in the EfR task, with no effect of chronic venlafaxine. A.*** *Escitalopram treated mice show no effect on number of chow bouts (p >0.05)* ***B.*** *Treated mice showed a reduction in the average time spent at chow bowl (p < 0.05), with no effect on* ***C.*** *latency to first time eating chow (p > 0.05) or* ***D.*** *latency to complete first trial (p > 0.05). Venlafaxine treated mice had no effect on* ***E.*** *number of chow bouts (p > 0.05).* ***F.*** *average time spent at chow bowl (p > 0.05),* ***G.*** *latency to first time eating chow (p > 0.05) or* ***H.*** *latency to complete first trial (p > 0.05). Error bars are mean ±SEM, unpaired t-test was performed to detect the drug effect, *p<0.05.*
